# Fibrinolytic niche is requested for alveolar type 2 cell-mediated alveologenesis and injury repair

**DOI:** 10.1101/2020.03.24.006270

**Authors:** Ali Gibran, Runzhen Zhao, Mo Zhang, Krishan G. Jain, Jianjun Chang, Satoshi Komatsu, Xiaohui Fang, Beiyun Zhou, Jiurong Liang, Dianhua Jiang, Mistuo Ikebe, Michael A Matthay, Hong-Long Ji

## Abstract

COVID-19, SARS, and MERS are featured by fibrinolytic dysfunction. To test the role of the fibrinolytic niche in the regeneration of alveolar epithelium, we compared the self-renewing capacity of alveolar epithelial type 2 (AT2) cells and its differentiation to AT1 cells between wild type (wt) and fibrinolytic niche deficient mice (*Plau^−/−^* and *Serpine1^Tg^*). A significant reduction in both proliferation and differentiation of deficient AT2 cells was observed *in vivo* and in 3D organoid cultures. This decrease was mainly restored by uPA derived A6 peptide, a binding fragment to CD44 receptors. The proliferative and differential rate of CD44^+^ AT2 cells was greater than that of CD44^−^ controls. There was a reduction in transepithelial ion transport in deficient monolayers compared to wt cells. Moreover, we found a marked suppression in total AT2 cells and CD44^+^ subpopulation in lungs from brain dead patients with acute respiratory distress syndrome (ARDS) and a mouse model infected by influenza viruses. Thus, we demonstrate that the fibrinolytic niche can regulate AT2-mediated homeostasis and regeneration via a novel uPA-A6-CD44^+^-ENaC cascade.

## BACKGROUND

The epithelial lining of regeneratively quiescent lungs is composed of alveolar type 2 (AT2) progenitor and differentiated alveolar type 1 (AT1) cells. To replace aged AT1 cells, AT2 cells undergo self-renewal to maintain alveolar epithelial homeostasis^1^. The regenerative potential of AT2 cells could be activated for timely recovery from lung epithelial injury^2–4^, including lobectomy and infections^5,6^. A marked suppression in fibrinolytic activity in local respiratory illnesses (*e.g*., inhaled smoke and aspirated gastric juice) and pulmonary complications of systemic diseases (*e.g*., sepsis) has been reported clinically and in animal models^7,8^. Migration and differentiation of mesenchymal stem cells (MSCs) in inflamed tissues are regulated by dynamic fibrinolytic niche^9–12^. The proteolysis of extracellular matrix substrates by urokinase plasminogen activator (uPA) could be involved in the benefit of the fibrinolytic niche to the regeneration of skeletal muscles and fractured cartilage^13–17^. For example, uPA and plasmin, two critical components of the fibrinolytic niche, cleave epithelial sodium channels (ENaC)^18,19^. In addition, functionally multifaceted uPA regulates alveolar epithelial function^20^ and possesses an A6 motif with a high affinity to CD44 receptors^21,22^. CD44^+^ AT2 cells show a higher proliferative capacity in fibrotic lungs^23^. The fibrinolytic niche in alveolar epithelial homeostasis and regeneration mediated by AT2 cells, however, has not been studied systematically. Our results have tested the potential novel contribution of uPA-PAI1-A6-CD44-ENaC cascade to the fibrinolytic niche in regulating the fate of AT2 cells.

## RESULTS

### Reduced proliferation and differentiation of AT2 cells in injured mouse and human lungs

To examine the effects of fibrinolytic niche on AT2 cells in normal and injured lungs, we infected wt and *Plau^−/−^* mice with the PR8 H1N1 type A influenza virus intranasally^24,25^. The severity of injured regions was classified as mild, moderate, and severe based on the lung injury score^26^. Randomly selected fields were analyzed with the ImageJ software to count proSPC^+^ AT2 cells (green) and the total cells for wt (**Fig. 1A**) and *Plau^−/−^* mice (**Fig. 1B**). Although AT1 cells were stained with anti-pdpn antibody, it was not possible to clearly distinguish from other non-epithelial cells, *i.e*., basal cells, leukocytes, endothelial cells, and fibroblasts due to considerable overlap. AT2 cells per field of wt lungs were approximately two-fold (12.6 ± 0.55%) that of *Plau^−/−^* lung preparations (6.0 ± 0.35 %, n=26 from 8 different mice). Moreover, influenza-infected lungs displayed widespread alveolar collapse and increased thickness of the lung interstitial septa in a severity-dependent manner. AT2 cells per field were significantly reduced in both moderate and severe injury regions as compared to wt controls (**Fig. 1A-B**). Similarly, this finding was observed in *Plau^−/−^* mice (**Fig. 1B-C**). AT2 cells per field for *Plau^−/−^* mice were markedly fewer than in wt animals (**Fig. 1C**. n=9 mice, 5 fields per mice). This is consistent with a reduction in the yield of total AT2 cells (1.7 × 10^6^ cells for *Plau^−/−^* mice vs 2.6 × 10^6^ cells for wt mice, **Fig. 1D**). We compared CD44^+^ AT2 cells between healthy and ARDS patients (n = 3 patients per group). As shown by fluorescence-activated cell sorting (FACS) data, CD44^+^ AT2 cells were reduced in ARDS patients (**Fig. 1E-G**). These results suggest that *Plau^−/−^* mice may mimic impaired fibrinolytic niches in the lung of ARDS patients.

**Figure 1.**
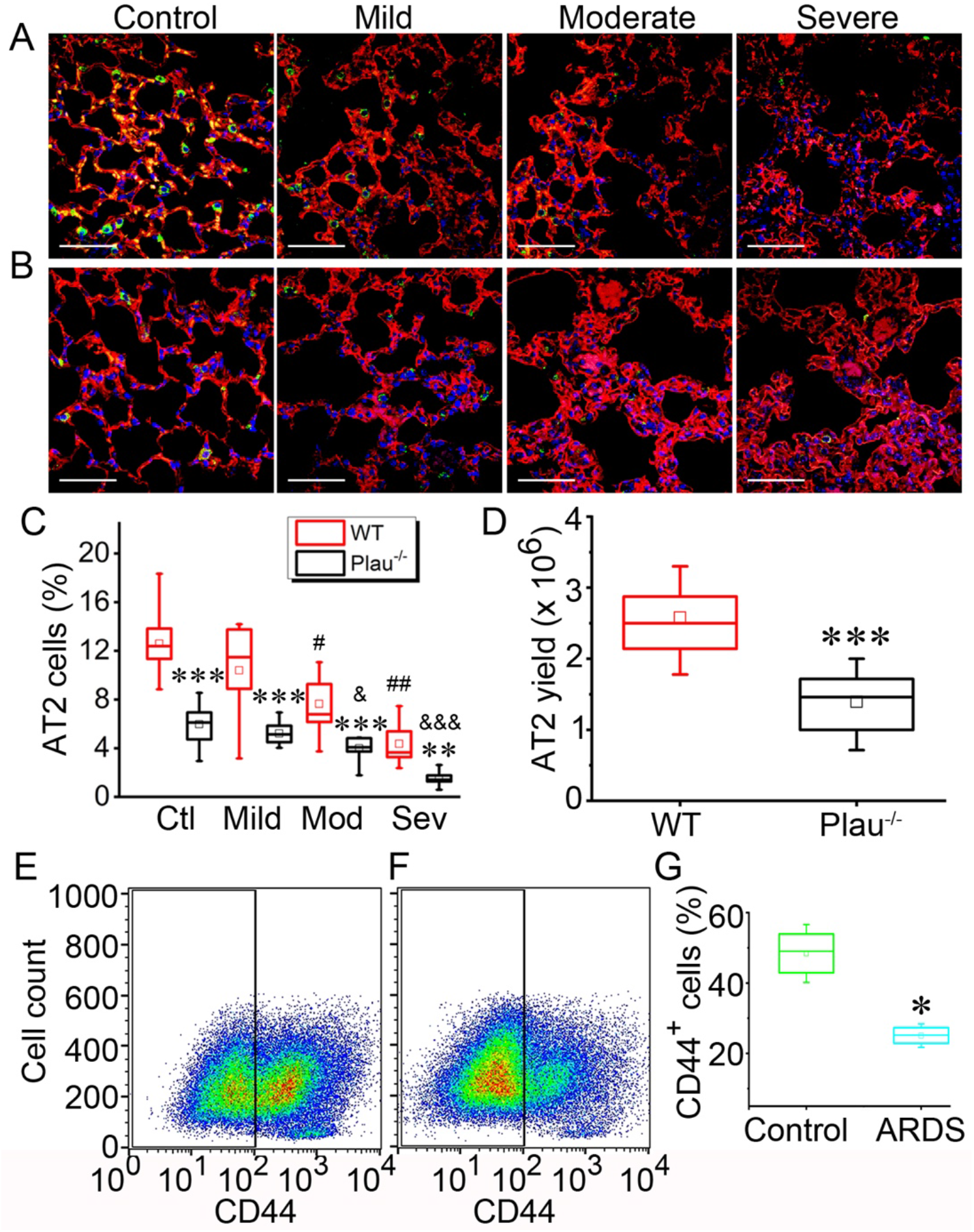
Quantification of AT2 cells in influenza-infected mouse lungs and ARDS lungs. Lung sections from control and influenza PR8 H1N1 infected mice were fixed and stained with anti-pdpn (AT1, red) and anti-sftpc (AT2, green) antibodies. Z-stacked confocal images of lung sections were prepared based on the severity of lung injury. (a) Images for wt mice 5 days post infection. n = 10 sections from 3 different mice per group. (b) Images for *Plau^−/−^* lungs. n = 10 sections from 7 different mice per group. (c) AT2 cells in normal and injured mouse lung sections. Mod, moderate. Sev, severe. n = 18 images from 3 different samples per group. (d-f) CD44^+^ AT2 cells isolated from ARDS lungs. (d) Yield of AT2 cells from *Plau^−/−^* and wt control mice. (n = 15 mice/genotype). Human AT2 cells isolated from 6 donors with FACS were slightly cultured for 24 – 48 h to improve viability, and then ran FACS for sorting CD44^+^ population was sorted by FACS for healthy control (e) and ARDS patients (f). (g) Average CD44^+^ cells (%). The data were analysed with one-way ANOVA followed by the Tukey *post hoc* test. n = 12 samples. * P < 0.05 vs healthy controls. Data in c and g are mean ± sem and analyzed with one-way ANOVA. ** P ≤ 0.01 and *** P ≤ 0.001 vs wt controls at the same severity. # P ≤ 0.05 and ## P ≤ 0.01 compared with controls for wt. & P ≤ 0.05 and &&& P ≤ 0.001 vs controls for *Plau^−/−^* group.

### *Plau* gene is required for AT2 mediated alveologenesis *in vitro*

To determine whether the *Plau* gene is required for re-alveolarization *in vitro*, we quantified spheroids formed by primary AT2 cells from both wt and *Plau^−/−^* mice in parallel (**Fig. 2A**). Organoids were captured as 4× DIC images from 4 to 12 days post seeding (**Fig. S2**). Organoids with a diameter > 50 μm (**Fig. 2B**) and colony forming efficiency (CFE) (**Fig. 2C**) were significantly decreased (n = 12, P < 0.01) in *Plau^−/−^* cultures compared with wt controls. Apparently, the organoids with a diameter ranging from 50 – 200 μm resulted in a reduction in colonies and CFE (**Fig. 2D**). The suppression in organoid formation associated with *Plau^−/−^* cells contributed to the reduction of total surface area, a clinical parameter for epithelial repair and development (**Fig. 2E**). The large organoids were filled with culture medium, having a smaller lumen with thicker walls for *Plau^−/−^* cultures over wt controls (**Fig. 2F-G**). These data provide direct evidence for the regulation of AT2-mediated re-alveologenesis by uPA.

**Figure 2.**
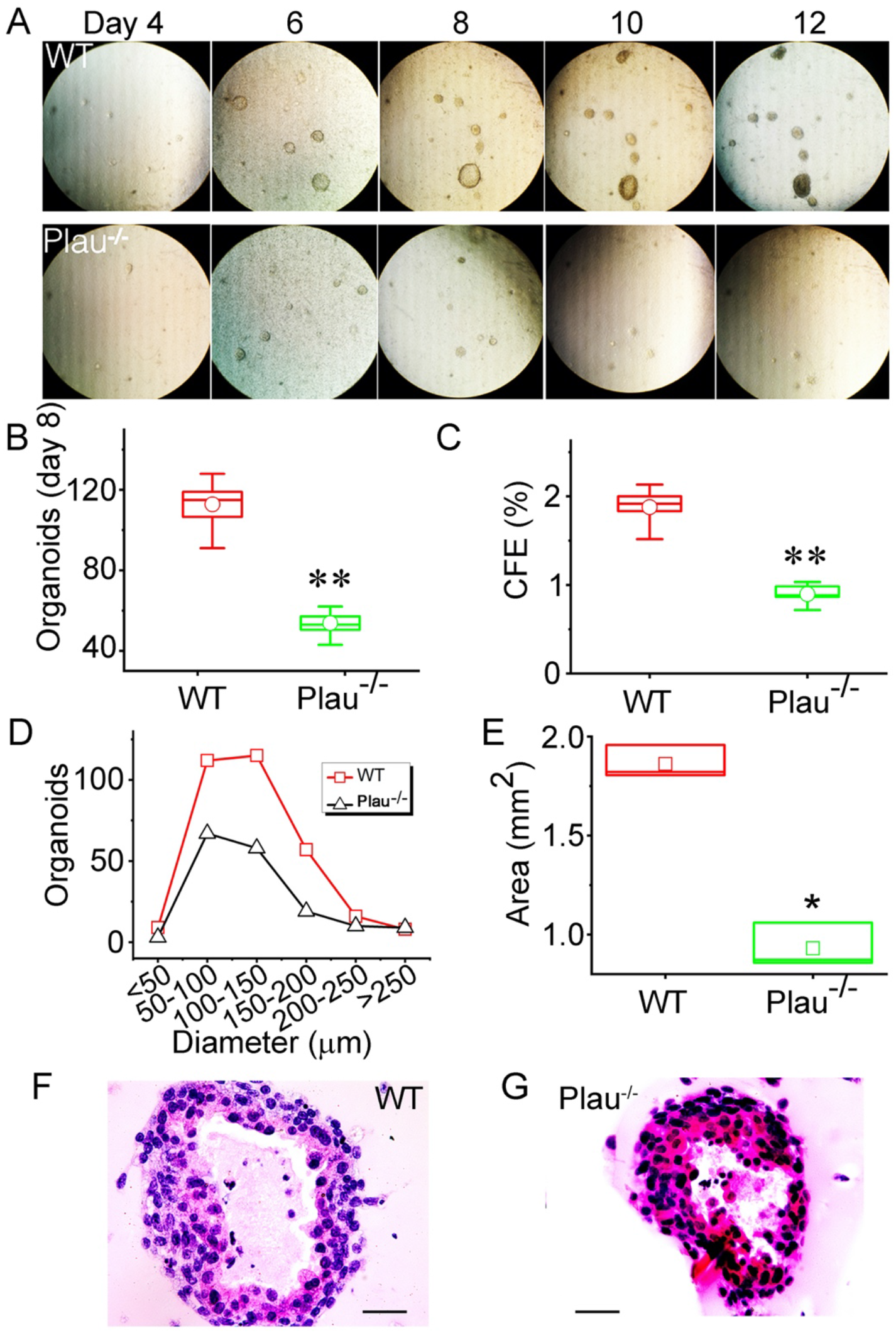
uPA is required for AT2 cells to develop spheroids in three-dimensional cultures. Colony formation observed and analyzed from day 4 to day 12 post seeding. (a) Representative phase-contrast images of colony per well for both wt and *Plau^−/−^* AT2 cells. n = 16 wells per set of experiments and repeated with 6 pairs of mice. (b-c) Colony number (b) and colony forming efficiency (CFE, c) on day 8 of the cultures. n=46. (d) Colony number as a function of the size (average diameter in μm). (e) Total organoid surface area per well. n = 46. Data in b, c, d, e, and g are mean ± sem and analyzed with the Student’s t-test. * indicates P ≤ 0.05 and ** P ≤ 0.01 compared with wt controls. (f-g) Representative images of organoid sections stained with the H & E procedure for wt (f) and *Plau^−/−^* group (g). Scale bar, 50 μm.

In addition, the organoids were further visualized with cell-permeable green fluorescent calcein to measure the cystic fibrosis transmembrane conductance regulator (CFTR) and ENaC activity (**Fig. S3**). The surface area was not altered in 30 minutes in the presence of cAMP-elevating forskolin, CFTRinh, and amiloride. One potential explanation is that these organoids do not develop tight junctions intercellularly as well as polarized monolayers.

### *Plau* gene regulates polarization and bioelectric features of AT2 monolayers

The divergent bioelectric features in polarized AT2 monolayers, including transepithelial resistance (R_T_) and short-circuit currents (I_SC_) were measured. *Plau^−/−^* monolayers showed a lower I_SC_ level compared to wt monolayer and diminished significantly by replacing Na^+^-free bath solution to inhibit Na^+^ ion transport (**Fig. 3A-B**), consistent with our previous studies in the airway epithelial cells^27^. Moreover, *Plau^−/−^* AT2 cells showed a reduced amiloride-sensitive I_SC_ level compared to wt cells (**Fig. 3C-D**). However, the amiloride affinity was not altered significantly (**Fig. 3D**). In addition, a greater R_TE_ value on day 5 was measured in wt monolayers compared to *Plau^−/−^* monolayer cultures (**Fig. 3E**). Thus, the ENaC activity seems to be regulated by the *Plau* gene.

**Figure 3.**
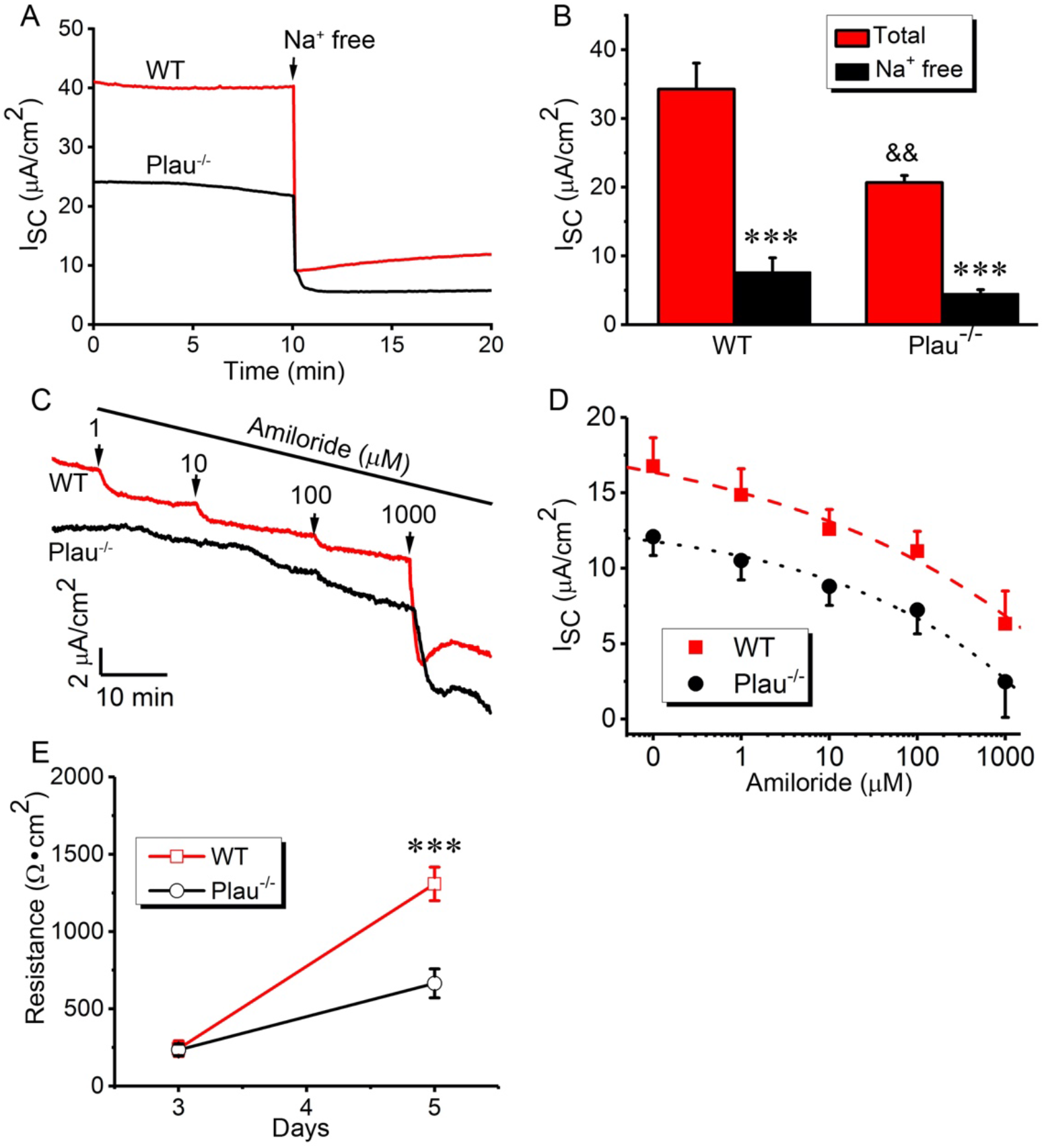
Bioelectric features of wt and *Plau^−/−^* AT2 monolayer cells. (a) Representative short-circuit current (Isc) in the presence and absence of bath Na^+^ ions. The bath solution was replaced with Na^+^ free medium as indicated by arrows. (b) Summarized Isc level. n=15. (c) Isc traces in the presence of accumulating amiloride added to the apical compartment. (d) Amiloride sensitivity of transepithelial Isc activity in AT2 monolayers. Dotted and dashed lines are generated by fitting raw data points with the Hill equation to compute IC50 value for amiloride. (e) Transepithelial resistance of AT2 monolayers measured with a chopstick meter. n = 15 monolayers from 3 different cell preparations. Data in b, d, and c are represented as mean ± sem and analyzed with the Student’s t-test. *** indicates P ≤ 0.001 vs basal Isc levels (b) or wt controls (e), && P ≤ 0.01 vs wt controls (b).

### *Plau* gene up-regulates the fate of AT2 cells

To quantify AT1 and AT2 cells in colonies, 3D images were obtained by stacking Z sections of each organoid (**Fig. S4**). Organotypic cultures showed a reduction in the number of epithelial cells in *Plau^−/−^* organoids as compared to wt cultures (**Fig. 4A-C**). Similarly, polarized monolayers exhibited a decline in both AT1 and AT2 cells in *Plau^−/−^* cells, too (**Fig. 4D-E**). Further, this reduction in AT1, AT2, and total cells in *Plau^−/−^* monolayers compared to wt cultures was confirmed by FACS (**Fig. 4F**). Also, cells counted in polarized monolayers showed a similar decline in both AT1 and AT2 cells (**Fig. S5A-B**). However, there was not a difference in the geometric volume and surface of monolayers between wt and *Plau^−/−^* cultures (**Fig. S5C-E**), perhaps due to the analysis of a small portion (0.12%, 0.0004 cm^2^) of entire monolayers (0.33 cm^2^). These results suggest that *Plau* regulates the renewal and differentiation of AT2 cells.

**Figure 4.**
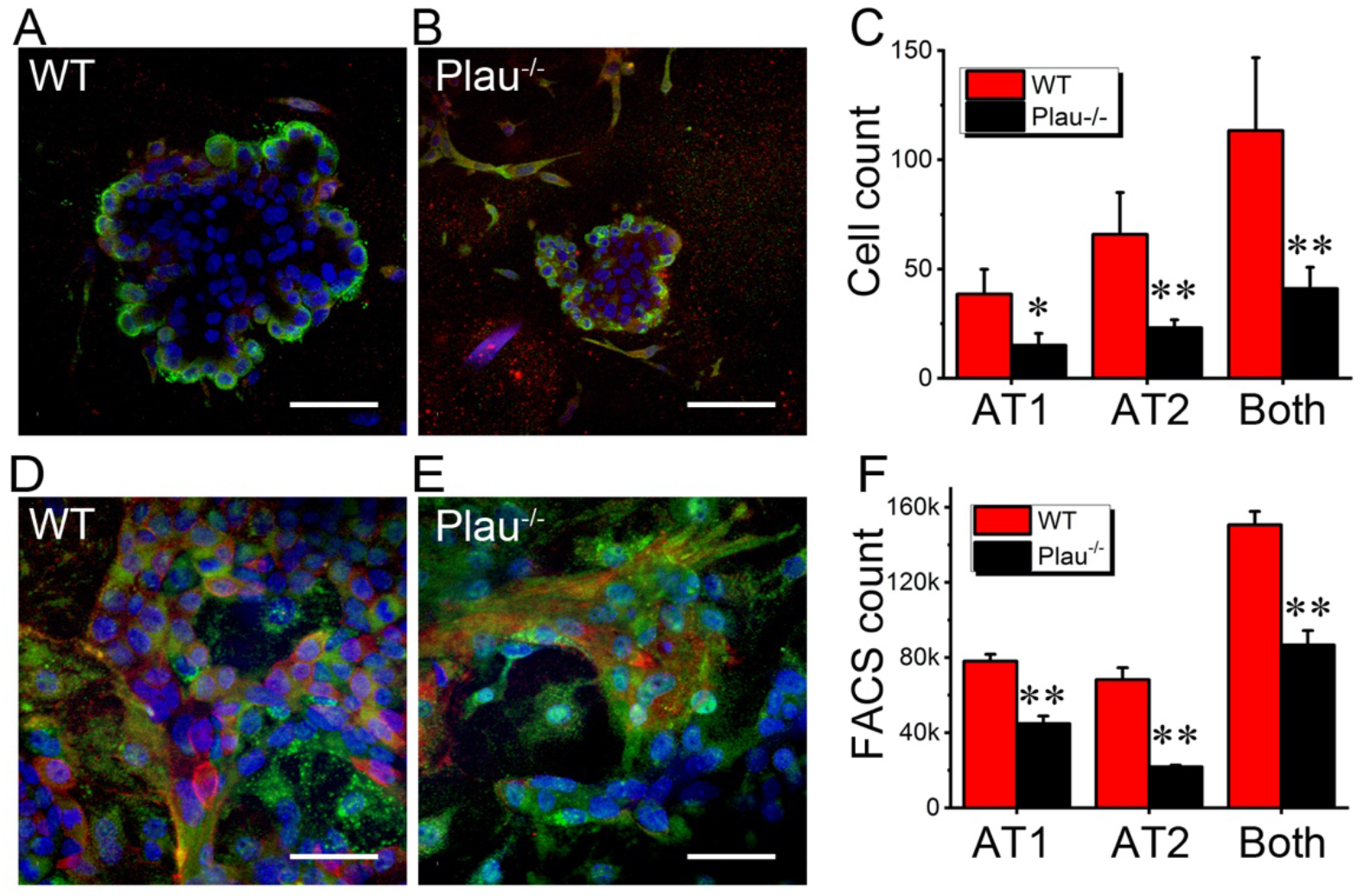
Proliferation and differentiation of AT2 cells in 3D organoid and polarized monolayer cultures. Cells were collected from organoids and monolayers on day 7 of the cultures at the air-liquid mode. (a-b) Stacked confocal images of wt (a) and *Plau^−/−^* organoids. AT1 and AT2 cells were identified as pdpn-positive (red) and sftpc-positive (green) cells, respectively. Stacked 3D image for each organoid was constructed, and cells were counted through the whole organoid. Scale bar, 100 μm. (c) Comparison of cell number for AT1, AT2 cells, and the sum of AT1 and AT2 cells (both). n = 16 organoids per group from three separate experiments. * P < 0.05 and ** P < 0.01 compared with wt. (d-e) Confocal images polarized AT2 monolayers. AT1 and AT2 biomarkers are pdpn and sftpc, respectively. Scale bar, 100 μm. (f) FACS analysis of cell types. n = 19 transwells per group from three separate experiments. ** P < 0.01 compared with wt. Data in c and f are means ±sem. The data were analyzed with the Student’s t-test.

### *Plau* gene facilitates DNA synthesis

To analyze DNA synthesis accompanied by cell proliferation, AT2 cells with active DNA synthesis were assessed by the EdU incorporation assays in both monolayers and organoids. Monolayer cultures of *Plau^−/−^* AT2 cells showed significantly lesser EdU^+^ cells (**Fig. 5A-B**). Further, organoids were analyzed with 3D stacking images to count EdU^+^ cells through an entire spheroid (**Fig. S6**). *Plau^−/−^* colonies showed a significantly reduced percentage of EdU^+^ cells compared with wt controls (**Fig. 5C-D**). The difference in EdU^+^ cells is in agreement with the diverse proliferative AT2 cells between wt and *Plau^−/−^* cultures.

**Figure 5.**
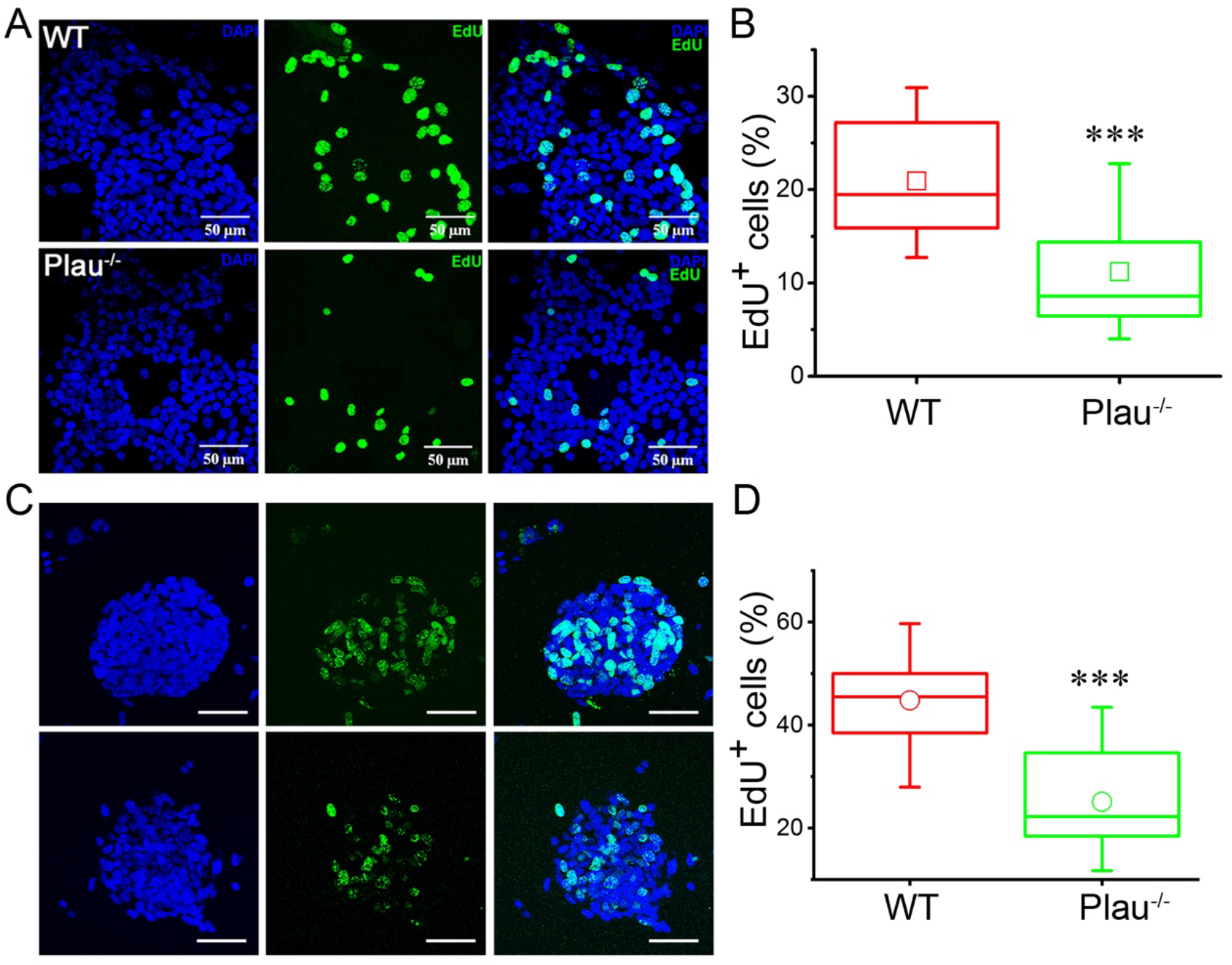
Detection of proliferative AT2 cells in organoids and polarized monolayers. A Click-iT EdU assay kit was used to identify AT2 cells with active DNA synthesis on day 7 of the cultures. (a) Images of wt (upper panel) and *Plau^−/−^* (lower panel) monolayers for DAPI (left), EdU (middle), and merged (right). Original magnification is 400 ×. Scale bar, 50 μm. (b) EdU^+^ cells (%) in wt and *Plau^−/−^* monolayers. n = 16 per group from 6 separate preparations. (c) Stacked organoid images of DAPI for nuclei (left), EdU (middle), and merged (right). Original magnification is 400 ×. Scale bar, 50 μm. (d) EdU^+^ cells (%) in wt and *Plau^−/−^* organoids. n = 25 for each group from 6 separate preparations. Data in b and d are means ± sem. The data were analyzed with the Student’s t-test. *** P ≤ 0.001 vs wt controls.

### The CD44 receptor and the uPA signal pathway involve in alveologenesis

The A6 peptide is derived from the connecting domain of uPA (from aa136 to aa143). The A6 peptide may serve as a mediator for uPA to bind with CD44 receptors located at the plasma membrane of AT2 cells. If so, the application of A6 peptides may restore the dysfunctional fate of *Plau* deficient AT2 cells. A6 peptides but not scramble controls (sA6) significantly increased spheroids (**Fig. 6A-B**). Also, A6 peptide markedly augmented the surface area of *Plau^−/−^* organoids per well (1.35 ± 0.13 mm^2^ vs. 0.82 ± 0.05 mm^2^ for sA6 group). In addition, blockade of CD44 receptors with a neutralizing antibody reduced wt organoids and corresponding surface area (**Fig. S6C**). Furthermore, the CD44 antibody resulted in a significant decrease in the number of AT2 cells in addition to a slight reduction in AT1 cells without statistical significance in wt organoids (**Fig. 6C, 6E-F**). In contrast, the number of both AT1 and AT2 cells was reduced in *Plau^−/−^* organoids (**Fig. 6C-F**), which were partially restored by A6 peptides. These two sets of experiments demonstrate that the binding of A6 peptides to CD44 receptors mediates the regulation of AT2 fate by uPA.

**Figure 6.**
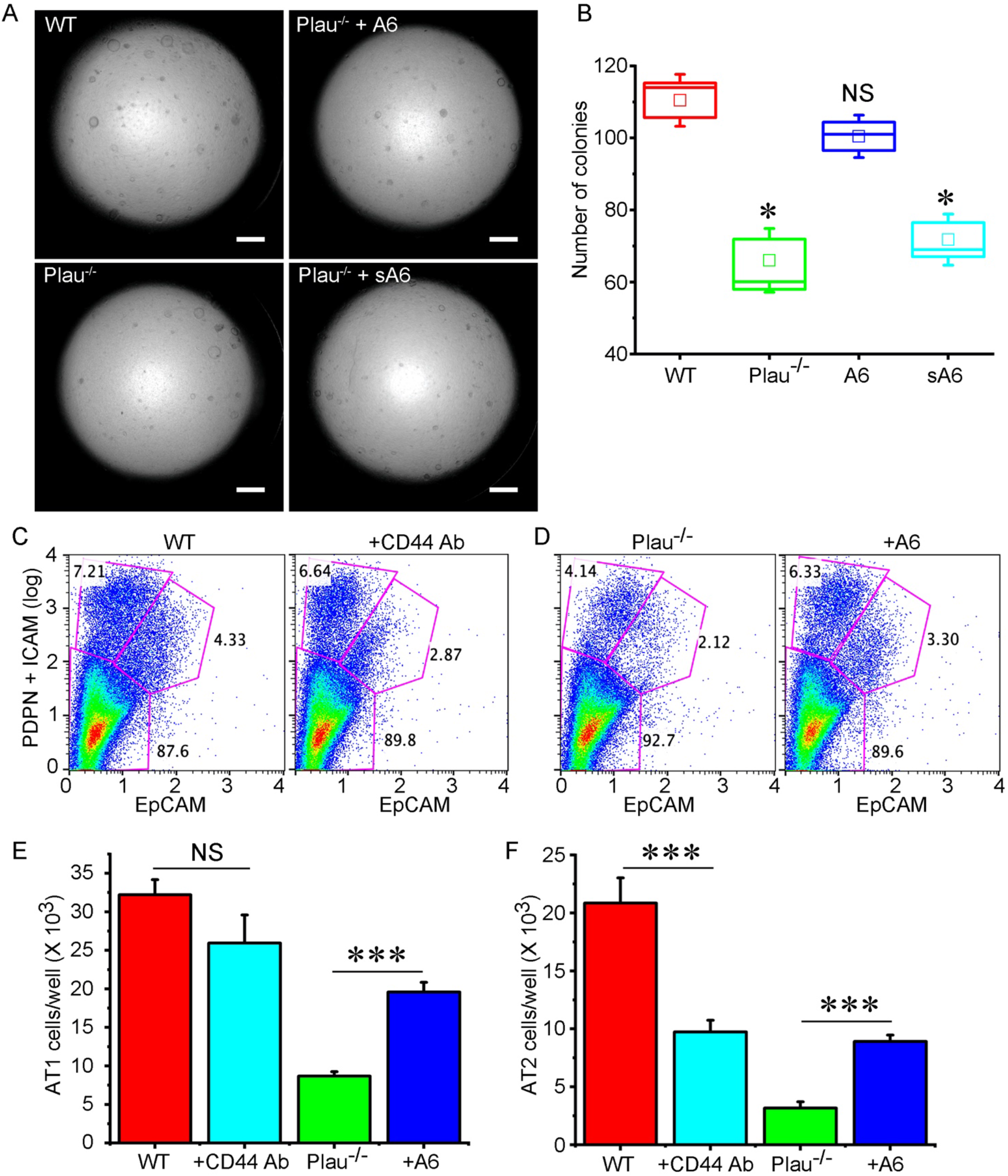
Effects of the A6 peptide and CD44 receptor on the formation of organoids and AT2 lineage. (a) Representative DIC images of organoids showing the effects of A6 peptides. AT2 cells isolated from *Plau^−/−^* mice were treated with either A6 derived from uPA or scrambled A6 peptides (sA6) in 3D organoid cultures with wt cultures as controls. (b) Organoid counts for each group. n = 16 transwells/ group from 6 independent experiments. (c-d) FlowJo analyzed FACS results. Wt AT2 cells were treated with blocking antibody against CD44 receptor (c), and *Plau^−/−^* cells were incubated with A6 peptide (d). Pdpn and EpCAM are biomarkers for AT1 and AT2 cells, respectively. (e-f) Cell sorting results for AT1 (d) and AT2 cells (f). n = 12 transwells for each group from 6 independent experiments. Data in b, e, and f are means ± sem, and analysed with one-way ANOVA followed by Tukey post hoc test. NS, not significant, * P ≤ 0.05, and *** P ≤ 0.001 compared with those groups as the horizontal lines indicated.

To corroborate these observations, we sorted and compared CD44^+^ and CD44^−^ AT2 cells between wt and *Plau^−/−^* mice by FACS. CD44 receptors at the AT2 cell surface were recovered for 24 – 36 h post enzymatic isolation. Significantly more CD44^+^ AT2 cells were harvested from wt mice than those from *Plau^−/−^* mice (**Fig. 7A-B**). We then cultured the organoids with these sorted CD44^+^ and CD44^−^ cells. Organoids developed by CD44^+^, *Plau^−/−^* AT2 cells were significantly reduced (112 ± 10 organoids vs. 182 ± 12 organoids for CD44^+^, *Plau*^+/+^ cells). In contrast, there was no significant difference in the number of organoids grown from CD44^−^ AT2 cells between wt and *Plau^−/−^* groups (30 ± 1 organoids for wt and 33 ± 2 organoids for *Plau^−/−^* line) (**Fig. 7C-E**). AT1 and AT2 cells in CD44^+^ *Plau^−/−^* organoids were fewer (n=6 transwells in 3 independent experiments) than wt CD44^+^ organoids (**Fig. 7F-G**). However, a significant decrease was observed only in AT1 but not AT2 cells of CD44^−^ *Plau^−/−^* organoids (**Fig. 7H**). Hence, both *in vivo* and *in vitro* data suggest that the *Plau* gene regulates the fate of CD44^+^ AT2 progenitor cells *in vivo* and that alveologenesis *in vitro* is utmost predominately determined by CD44^+^ cells.

**Figure 7.**
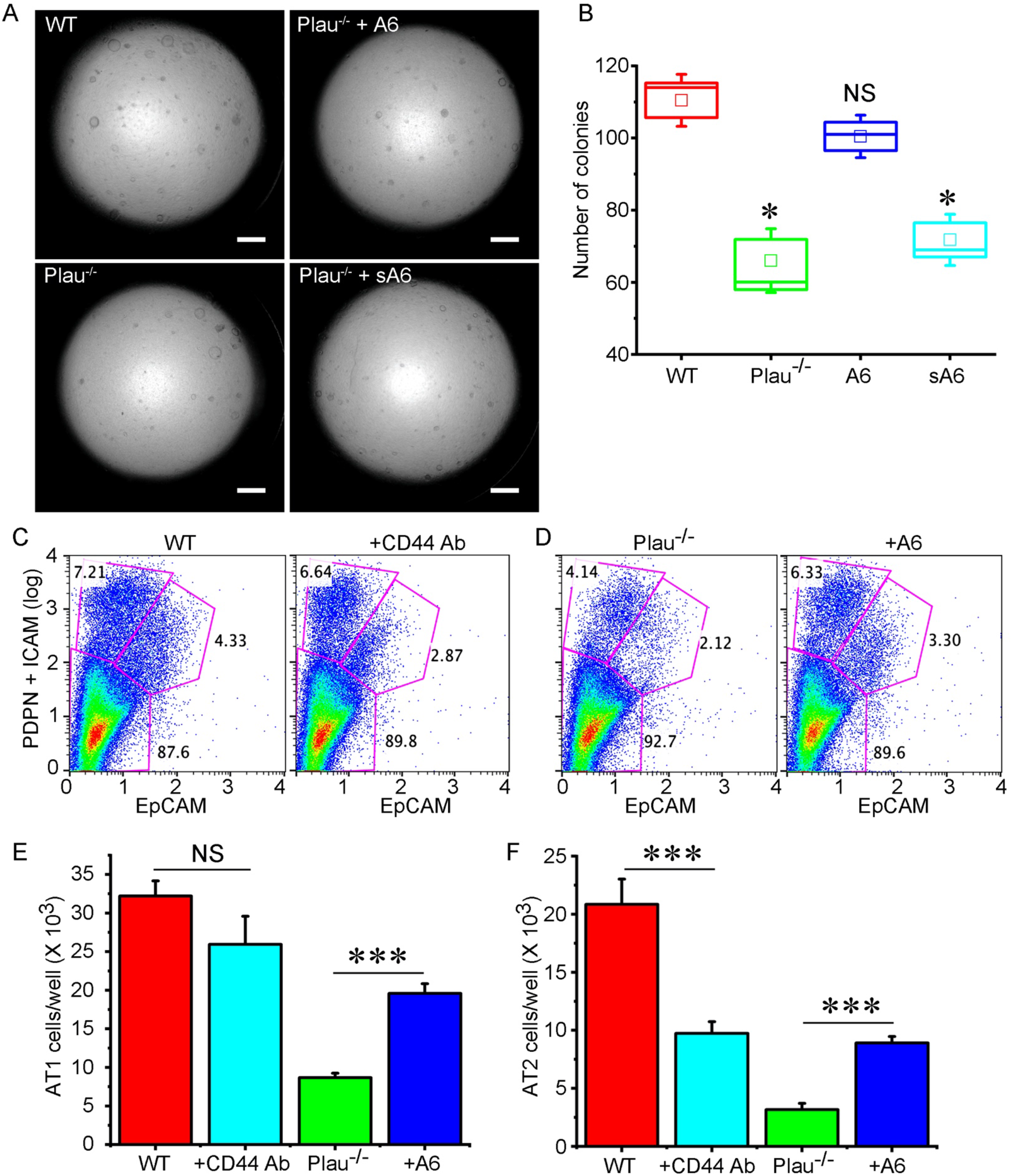
Roles of CD44 expression in AT2-mediated alveologenesis *in vitro*. Potentially impaired CD44 receptors at the plasma membrane of AT2 cells were restored by culturing on collagen IV coated plates cultured for 24 – 36 h at 37°C. Then cells were detached and sorted as EpCAM^+^CD44^+^ and EpCAM^+^CD44^−^ subpopulations. (a) FACS sorted EpCAM^+^CD44^+^ AT2 cells for wt (left) and *Plau*^−/−^ (right) groups. (b) Comparison of the number of EpCAM^+^CD44^+^AT2 cells between wt and *Plau^−/−^* mice. n = 7 mice / genotype. (c) DIC images of organoids formed by sorted CD44^+^ (upper panel) and CD44^−^ (lower panel) AT2 cells on day 7. (d-e) Number of organoids per transwell for CD44^+^ (d) and CD44^−^ (e) AT2 cells. n = 18 for each group from four independent experiments. (f) FACS analysis of AT1 (PdPn^+^ICAM^+^) and AT2 (EpCAM^+^) cells. (g-h) Statistical comparison of AT1 and AT2 cells in organoids grown from CD44^+^ (g) and CD44^−^ (h) subpopulations. n = 12 transwells per group from three independent experiments. Data in b, d, e, g, and h are means ± sem and analysed with the Student’s t-test in b, pair signed rank test in d-e and one-way ANOVA followed by Tukey post hoc analysis in g-h. NS, not significant, * P ≤ 0.05, and *** P ≤ 0.001 compared with wt controls.

### Fibrinolytic activity is required for re-alveolarization and tissue homeostasis

To further substantiate the role of the fibrinolytic niche in lung regeneration, we compared the proliferation and differentiation of AT2 cells between wt and *Serpine1* transgenic (*PAI-1^Tg^*) mice. Elevated PAI-1 level is a hallmark of lung injury patients^28–33^. Genetically engineered humanized mice expressing a gain-of-function PAI-1 human gene mimicked the disrupted fibrinolytic niche in septic ARDS patients^34,35^. Lesser AT2 cells were harvested from *PAI-1^Tg^* mice compared to wt (**Fig. 8A**). These cells were then sorted for CD44^+^ and CD44^−^ cells (**Fig. 8B**). A significant decrease in CD44^+^ AT2 cells was observed in *PAI-1^Tg^* mice, which was similar to that in *Plau^−/−^* mice (**Fig. 8C**). Moreover, CD44^+^ AT2 cells of *PAI-1^Tg^* mice developed fewer organoids than wt controls (61 ± 9 organoids vs. 91 ± 7 organoids for wt controls, n=12) (**Fig. 8D-E**). Correspondingly, the total surface area of organoids of *PAI-1^Tg^* AT2 cells was significantly reduced compared with wt cultures (**Fig. 8F**). Further, duplication of *PAI-1^Tg^* AT2 but not AT1 cells in organoids were down-regulated (**Fig. 8G-H**). On the other hand, the transepithelial resistance in wt AT2 monolayers was reduced with a maximal difference on day 5 than that in *PAI-1^Tg^* cultures (**Fig. 8I**). Basal, amiloride-sensitive, and amiloride-resistant fractions of the short-circuit (I_SC_) currents in *PAI-1^Tg^* monolayers were reduced (**Fig. 8J**). However, the amiloride sensitivity remained unchanged by over-expression of the PAI-1 gene, as shown by apparent *k_i_* values (**Fig. 8K**). Consistent with the results in *Plau^−/−^* mice and cultures, disruption of the fibrinolytic niche by PAI-1 overexpression attenuates the rate of AT2 cells for renewal and differentiation to AT1 cells.

**Figure 8.**
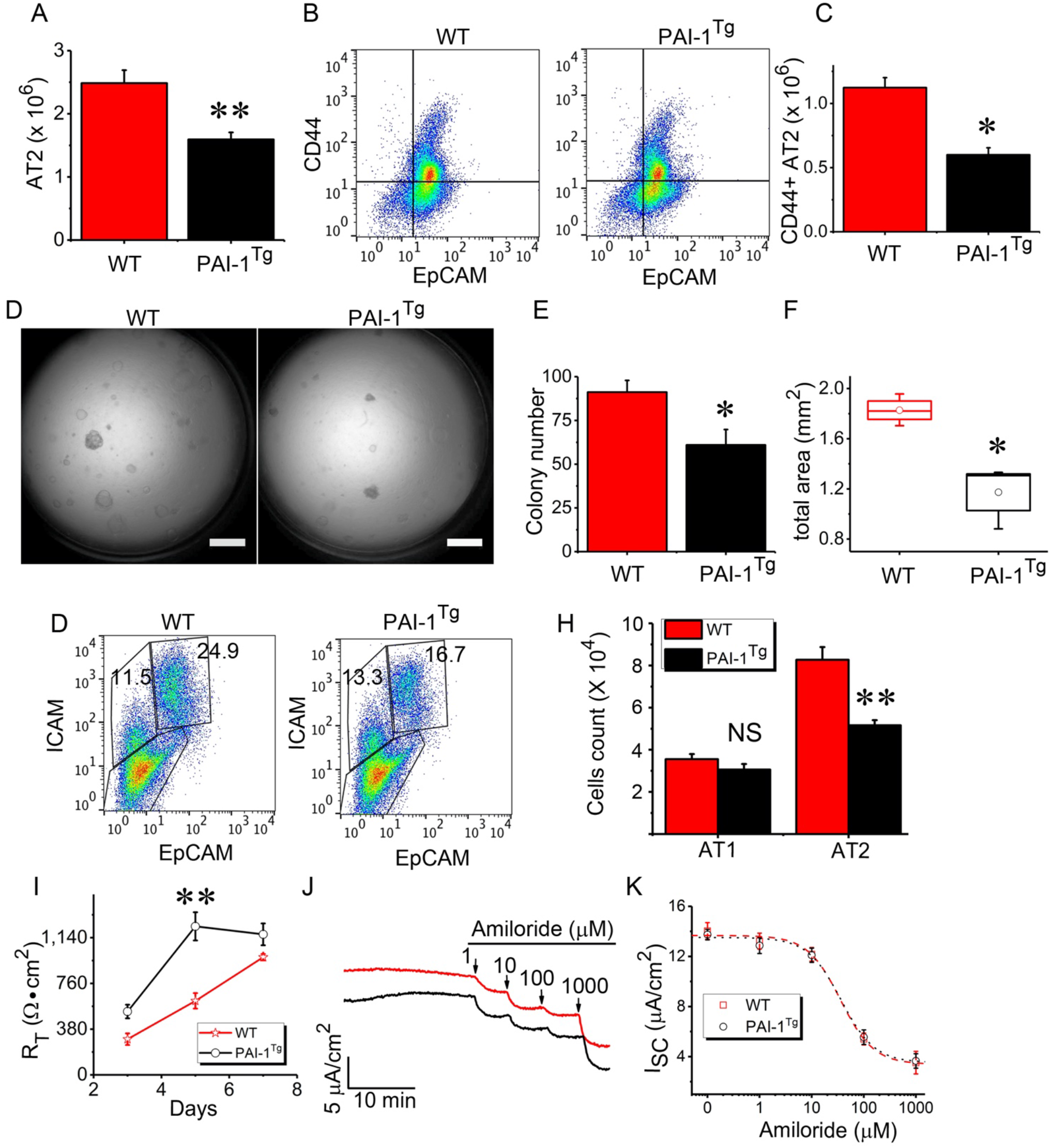
Effects of PAI-1 overexpression on the regenerative and channel function of AT2 cells. AT2 cells were isolated from wt and *PAI-1^Tg^* mice overexpressing mutated gain-of-function human PAI-1 gene. (a) Different yields of AT2 cells between wt and *PAI-1^Tg^* mice. n = 11 pairs of mice. AT2 cells were harvested in parallel from one pair of wt and *PAI-1^Tg^* mice per experiment. (b) Comparison of the yield of FACS-sorted CD44^+^ AT2 cells between wt and *PAI-1^Tg^* mice. (c) Difference in the number of CD44^+^ AT2 cells *in vivo* between wt and *PAI-1^Tg^* mice. n = 9 pairs of mice. (d) DIC images of AT2 organoids for wt (left) and *PAI-1^Tg^* mice. Scale bar, 100 μm. (e-f) Organoid number (e) and total surface area (f) of organoids per transwell. n = 15 transwells per group. (g) FACS analysis of AT1 (ICAM^+^) and AT2 (EpCAM^+^) cells. (h) Cell count for AT1 and AT2 cells. n = 3 pairs of mice. (i) Transepithelial resistance (R_T_) of AT2 monolayers measured with a chopstick meter. n = 16 monolayers per group. (j) Representative short-circuit current (Isc) traces in response to accumulating amiloride doses. (k) Amiloride sensitivity. Raw data from (j) were fitted with the Hill equation to compute IC50 value. n = 12. Data in a, c, e, f, h, i, and k are mean ± s.e.m. and analysed with one-way ANOVA followed by the Tukey post hoc test in (i) and the Student’s t-test in others. NS, not significant, * P ≤ 0.05, and ** P ≤ 0.01 compared with wt controls.

## DISCUSSION

The primary objective of this study was to decipher the role of the fibrinolytic niche in the re-alveolarization mediated by progenitor AT2 cells and underlying mechanisms in healthy and injured lungs. We employed influenza-infected mice, *Plau*^−/−^ and *Serpine1^Tg^* mouse strains, primary human AT2 cells from the human lungs of brain-dead patients with ARDS, and 3D organoids and polarized monolayers of AT2 cells to trace the cell fate *in vivo* and *in vitro*. The results demonstrate that the impaired fibrinolytic niche in influenza-infected lungs and genetically engineered mice modeling impaired fibrinolytic niche results in a significant decline in the proliferative AT2 and differentiated AT1 cells.

These results thus demonstrate the novel finding that that the fibrinolytic niche may be critical for re-alveologenesis. Previous studies suggest that *Plau^−/−^* and *PAI-1^Tg^* mice are more susceptible to lung fibrosis post bleomycin injury^36,37^. Clinically, impaired fibrinolytic niche, including elevated PAI-1 contents and suppressed fibrinolytic activity, are prognostic markers for ARDS patients^38,39^. Deficiency of plasminogen activators and plasminogen leads to dysfunctional repair of injured tissues (*i.e*., skin and muscle)^40,41^. In contrast, regeneration of injured muscle is improved by knocking out the *Serpine1* gene1^42^. The *Plau* gene regulates the proliferation of various cells, including renal epithelial cells^43,44^, umbilical vein endothelial cells^45^, vascular endothelial cells^46^, and keratinocytes^47,48^. Contrary to this, *Serpine1* is a negative regulator of cell proliferation through the phosphatidylinositol3-kinase/Akt pathway in endothelial cells^49,50^. Our study uncovers a new signaling pathway of the fibrinolytic niche in the AT2 cell-mediated re-alveolarization. The uPA/A6/CD44/ENaC cascade is involved in the regulation of self-renewal and differentiation of mouse and human AT2 cells, particularly the CD44^+^ subpopulation. This conclusion was further substantiated in *Serpine1^Tg^* mice that have a disrupted fibrinolytic niche similar to the *Plau^−/−^* mice, and both mouse lines are reasonable pre-clinical models of ARDS. The proteolytic link between the fibrinolytic niche and ENaC proteins has been provided by other groups and us *in vivo* and *in vitro*^27,51–53^. The differences between each component of the fibrinolytic niche are described previously. For example, uPA but not tPA cleaves human gENaC subunits^51^. PAI-1 or antiplasmin alone does not affect ENaC activity. The slight diversity between the *Plau^−/−^* and *Serpine1^Tg^* models in this study could be due to their differential regulation of ENaC and other down-stream molecules.

CD44 is a crucial mediator for the regulation of re-alveologenesis by AT2 cells. We previously demonstrated that CD44 receptors govern cell survival and the progression of lung fibrosis via the Toll-like receptors and hyaluronan^54^. The connective domain of uPA has an eight L-amino acid sequence (Ac-KPSSPPEE-NH2), known as A6 peptide^55^. Binding of the A6 peptide with the link domain of CD44 receptors initiates the downstream signaling^56^. *CD44^−/−^* mice demonstrate an inflammatory phenotype characterized by increased inflammatory cell recruitment and are more susceptible to LPS induced injury. *CD44* deficiency leads to impaired expression of negative regulators of TLR signaling in macrophages, an essential event in the prevention of LPS induced inflammatory responses^57^. Our study demonstrates a new mechanism that the CD44 receptors at the plasma membrane of AT2 cells are a key player for AT2 cell-mediated re-alveolarization. In addition to the reduction in AT2 cells in injured mouse and human lungs associated with aberrant fibrinolytic niche, both *Plau^−/−^* and *Serpine1^Tg^* AT2 cells cannot develop organoids equal to those of wt mice *in vitro*, mostly likely due to the lesser expression of CD44 receptors in AT2 cells as A6 peptide compensates for the loss of organoids. This is supported by our previously unreported observations that CD44^+^ AT2 cells have an enhanced capacity for selfrenewal and differentiation to AT1 cells. Of note, both *Plau^−/−^* and *Serpine1^Tg^* mice have lesser CD44^+^ AT2 cells so that the total yield of primary AT2 cells is significantly reduced compared with wt controls.

The brain ENaC proteins encoded by the *Scnn1a, 1b, 1d*, and *1g* are involved in adult neurogenesis by the transmembrane sodium signals for the deletion of *Scnn1a* or functional blockade by amiloride in neural stem cells strongly impairs their proliferation and differentiation^58^. ENaC has been functionally detected in both club cells and AT2 cells^59^. Whether ENaC is a member of the fibrinolytic niche for lung stem cell-mediated regeneration is unknown. We recently reported that overexpression of the *Scnn1d*, a pseudogene in mice, significantly improves the development of organoids in 3D cultures of primary mouse AT2 cells^60^. Together with abnormal ENaC activity in *Plau^−/−^* and *Serpine1^Tg^* AT2 cells, we conclude that the ENaC signal may be an essential regulator for the fate of AT2 cells in normal and injured lungs.

In summary, our study uncovers a novel role of the fibrinolytic niche in alveolarization via the uPA/A6/CD44 and the uPA/ENaC signaling pathways. This study provides new evidence for the possible prognostic relevance of the fibrinolytic niche for ARDS patients. Targeting the abnormality of the fibrinolytic niche could be a promising pharmaceutic strategy to accelerate reparative processes of injured alveolar epithelium in clinical disorders such as ARDS.

## METHODS

Methods, including statements of data availability and any associated accession codes and references, are available in the online version of this paper.

## ACKNOWLEDGMENTS

This work was supported by NIH grant R01 HL134828 and Donor West (California).

## AUTHOR CONTRIBUTIONS

Conceptualization: H.L.J. and M.A.M; investigation: G.A., R.Z.Z, J.C., M.Z., S.K., X. F., B. Z., J.L., D. J. M. I., K.G.J.; formal analysis: G.A, H.L.J, D. J., K.G.J.; writing: G.A., H.L.J., M.A.M.; supervision: H.L.J., M.A.M.; funding: H.L.J., M.A.M.

## Supplementary file

### MATERIALS & METHODS

#### Cells, antibodies, reagents, and key materials

Major critical reagents are listed in the table. General chemicals and reagents were purchased from Sigma.Supplementary file

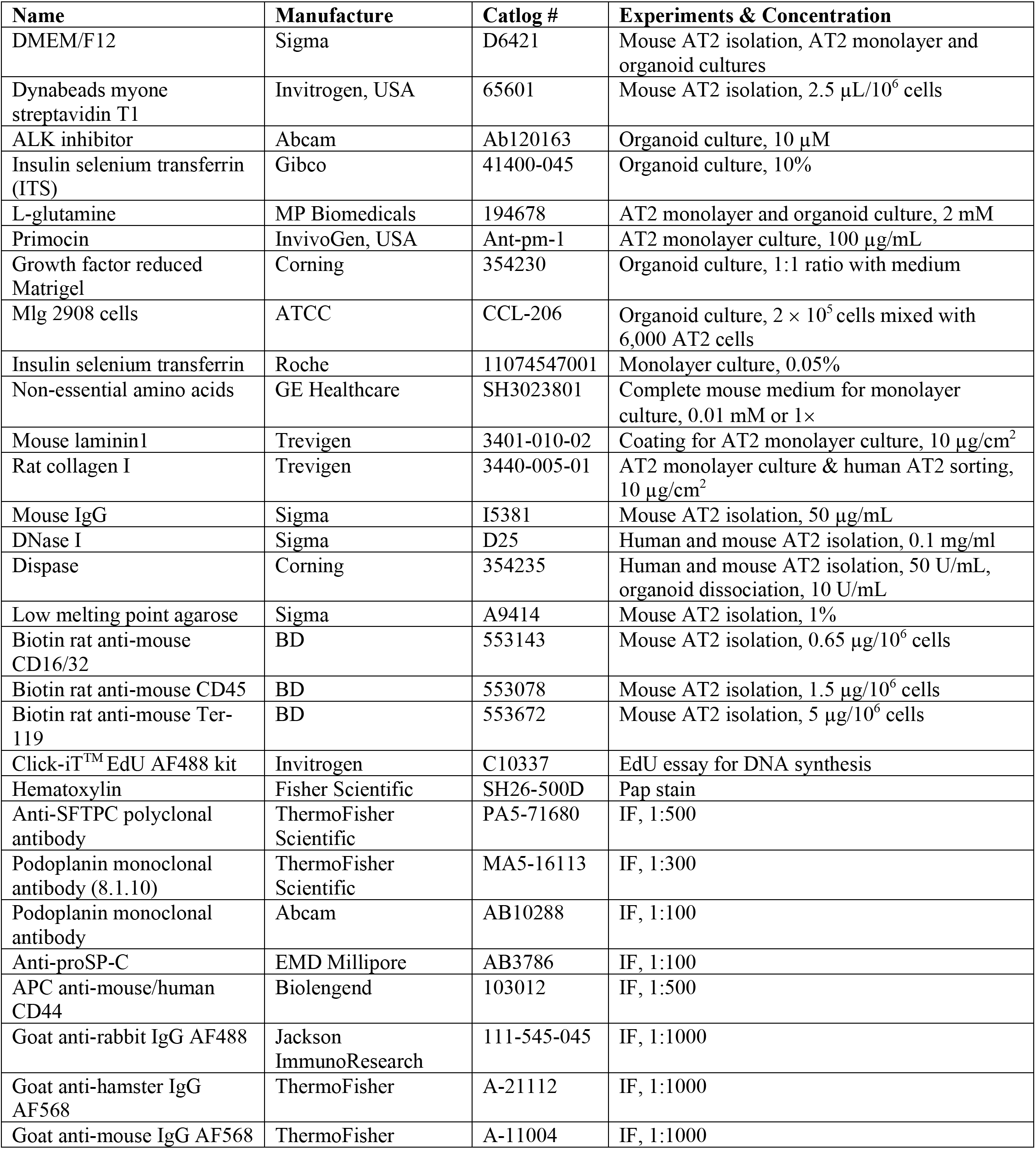
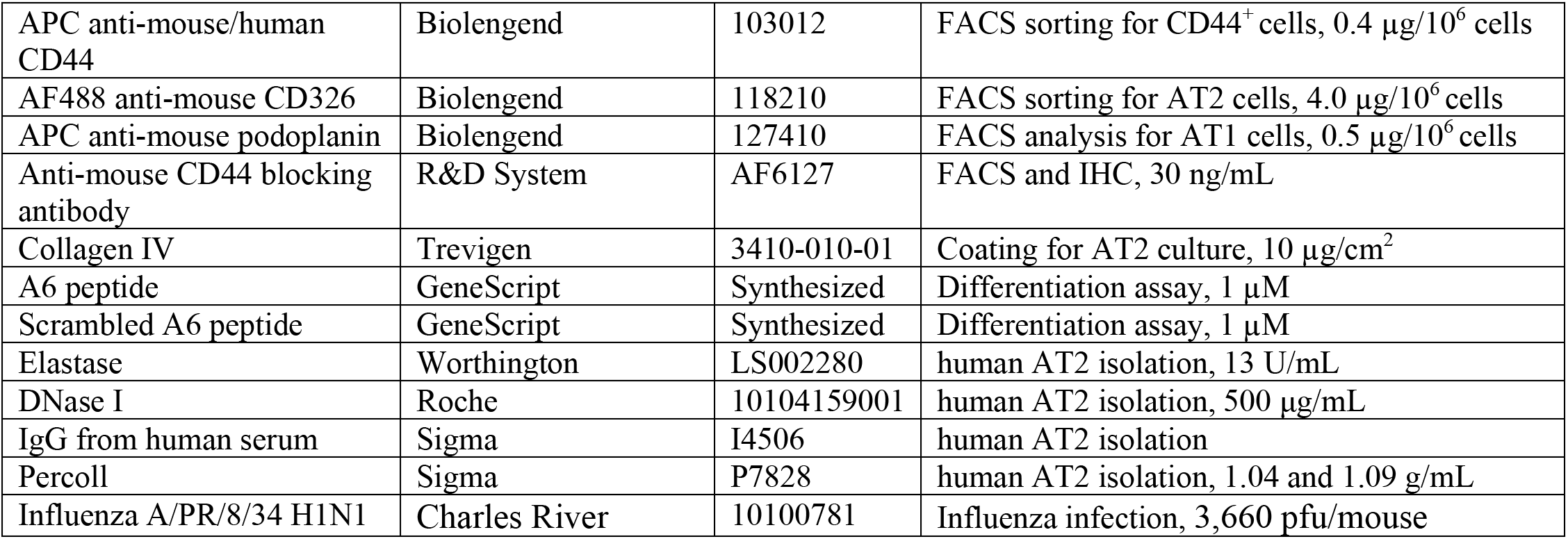

| **Name** | **Manufacture** | **Catlog #** | **Experiments & Concentration** |
| --- | --- | --- | --- |
| DMEM/F12 | Sigma | D6421 | Mouse AT2 isolation, AT2 monolayer and organoid cultures |
| Dynabeads myone streptavidin T1 | Invitrogen, USA | 65601 | Mouse AT2 isolation, 2.5 μL/$10^6$ cells |
| ALK inhibitor | Abcam | Ab120163 | Organoid culture, 10 μM |
| Insulin selenium transferrin (ITS) | Gibco | 41400-045 | Organoid culture, 10% |
| L-glutamine | MP Biomedicals | 194678 | AT2 monolayer and organoid culture, 2 mM |
| Primocin | InvivoGen, USA | Ant-pm-1 | AT2 monolayer culture, 100 μg/mL |
| Growth factor reduced Matrigel | Corning | 354230 | Organoid culture, 1:1 ratio with medium |
| Mlg 2908 cells | ATCC | CCL-206 | Organoid culture, $2 \times 10^5$ cells mixed with 6,000 AT2 cells |
| Insulin selenium transferrin | Roche | 11074547001 | Monolayer culture, 0.05% |
| Non-essential amino acids | GE Healthcare | SH3023801 | Complete mouse medium for monolayer culture, 0.01 mM or 1× |
| Mouse laminin1 | Trevigen | 3401-010-02 | Coating for AT2 monolayer culture, 10 μg/cm<sup>2</sup> |
| Rat collagen I | Trevigen | 3440-005-01 | AT2 monolayer culture & human AT2 sorting, 10 μg/cm<sup>2</sup> |
| Mouse IgG | Sigma | I5381 | Mouse AT2 isolation, 50 μg/mL |
| DNase I | Sigma | D25 | Human and mouse AT2 isolation, 0.1 mg/ml |
| Dispase | Corning | 354235 | Human and mouse AT2 isolation, 50 U/mL, organoid dissociation, 10 U/mL |
| Low melting point agarose | Sigma | A9414 | Mouse AT2 isolation, 1% |
| Biotin rat anti-mouse CD16/32 | BD | 553143 | Mouse AT2 isolation, 0.65 μg/$10^6$ cells |
| Biotin rat anti-mouse CD45 | BD | 553078 | Mouse AT2 isolation, 1.5 μg/$10^6$ cells |
| Biotin rat anti-mouse Ter-119 | BD | 553672 | Mouse AT2 isolation, 5 μg/$10^6$ cells |
| Click-iT™ EdU AF488 kit | Invitrogen | C10337 | EdU essay for DNA synthesis |
| Hematoxylin | Fisher Scientific | SH26-500D | Pap stain |
| Anti-SFTPC polyclonal antibody | ThermoFisher Scientific | PA5-71680 | IF, 1:500 |
| Podoplanin monoclonal antibody (8.1.10) | ThermoFisher Scientific | MA5-16113 | IF, 1:300 |
| Podoplanin monoclonal antibody | Abcam | AB10288 | IF, 1:100 |
| Anti-proSP-C | EMD Millipore | AB3786 | IF, 1:100 |
| APC anti-mouse/human CD44 | Biolengend | 103012 | IF, 1:500 |
| Goat anti-rabbit IgG AF488 | Jackson ImmunoResearch | 111-545-045 | IF, 1:1000 |
| Goat anti-hamster IgG AF568 | ThermoFisher | A-21112 | IF, 1:1000 |
| Goat anti-mouse IgG AF568 | ThermoFisher | A-11004 | IF, 1:1000 |
| APC anti-mouse/human CD44 | Biolengend | 103012 | FACS sorting for CD44<sup>+</sup> cells, 0.4 µg/10<sup>6</sup> cells |
| AF488 anti-mouse CD326 | Biolengend | 118210 | FACS sorting for AT2 cells, 4.0 µg/10<sup>6</sup> cells |
| APC anti-mouse podoplanin | Biolengend | 127410 | FACS analysis for AT1 cells, 0.5 µg/10<sup>6</sup> cells |
| Anti-mouse CD44 blocking antibody | R&D System | AF6127 | FACS and IHC, 30 ng/mL |
| Collagen IV | Trevigen | 3410-010-01 | Coating for AT2 culture, 10 µg/cm<sup>2</sup> |
| A6 peptide | GeneScript | Synthesized | Differentiation assay, 1 µM |
| Scrambled A6 peptide | GeneScript | Synthesized | Differentiation assay, 1 µM |
| Elastase | Worthington | LS002280 | human AT2 isolation, 13 U/mL |
| DNase I | Roche | 10104159001 | human AT2 isolation, 500 µg/mL |
| IgG from human serum | Sigma | I4506 | human AT2 isolation |
| Percoll | Sigma | P7828 | human AT2 isolation, 1.04 and 1.09 g/mL |
| Influenza A/PR/8/34 H1N1 | Charles River | 10100781 | Influenza infection, 3,660 pfu/mouse |

#### Animal husbandry

All mice purchased from Jackson Laboratory were maintained in a pathogen-free facility. A 12-h light/dark cycle and ad libitum supply for food and water were provided. Age, sex, and weight-matched (4-12 months) wild type (wt), *Plau^−/−^*, and *Sepine1^Tg^* mice were sacrificed for experiments as approved by the Institute of Animal Care and Use Committee of the University of Texas Health Science Center at Tyler.

#### Influenza-induced lung injury and immunohistology

Wt and *Plau^−/−^* mice were infected intranasally with a dose of 3,660 pfu influenza virus (type A/PR8/34 H1N1, Charles River, USA) diluted in 50 μl PBS per anesthetized animal. Control animals were delivered intranasally with the same amount of PBS. Both control and infected mice were euthanized 5 d post infection. The trachea was tied with a suture followed by opening the chest cavity and removing the lungs subsequently. The dissected lungs were fixed with 10% neutral buffered formalin v/v (Richard-Allan Scientific) for 72 h at room temperature. Lung tissues were then dehydrated with increasing ethanol grades, embedded in paraffin, and sectioned at a thickness of 7 μm.

#### Visualization and quantification of AT2 cells in mouse and human lung tissues

Lung sections from both influenza-infected mice and brain dead patients who had ARDS based on the Berlin criteria were deparaffinized and rehydrated in xylene and a series of ethanol at cumulating concentrations and in H_2_O for 5 min. To unmask antigens, tissue slides were incubated with 10 mM sodium citrate buffer (Thermo Scientific), pH 6.0, for 20 min at 95°C, cooled for another 20 min at room temperature, and then washed with PBS. After blocking for 1 h with 1% BSA and 4% normal goat serum, tissue sections were incubated with following antibodies: anti-proSP-C (1:500, EMD Millipore), anti-pdpn (1:300, ThermoFisher) for mouse lung tissues and 1:100 dilution for both antibodies (Abcam) for human lung slices. After 3 times washing with PBS, secondary antibodies were applied: goat anti-rabbit IgG AF488, goat anti-hamster AF568, and goat anti-mouse AF568. Tissue sections were stained with DAPI and sealed. Images were captured using a Zeiss LSM 510 confocal microscope. Images were captured containing at least 7 – 8 individual 1μm optical Z-sections. Z-stacks were obtained from at least five randomly selected areas. All images were subsequently processed with ImageJ software. AT2 cells were counted using a cell counter plug-in for ImageJ software. Cells were counted in at least 5 different areas to obtain a total above 1,000 cells and then analyzed statistically.

#### Mouse AT2 isolation

Mouse AT2 cells were isolated from wt, *Plau^−/−^*, and *Sepine1^Tg^* strains of C57BL/6 animals (Jackson Laboratory, USA) as previously reported with modifications^1^. Briefly, mice were euthanized and exsanguinated, followed by perfusing lungs with 10 – 20 mL DPBS until pink lungs turned to white. The trachea was cannulated with a 20G catheter to instill 1.5 – 2.0 mL dispase followed by 0.5 mL of 1% low melting point agarose. The lungs were dissected and incubated in 50 U/mL dispase solution for 45 min at room temperature. The lungs were gently teased in DMEM/F-12 + 0.01% DNase I and incubated for 10 min at room temperature. Cells were passed through a serial filtration (100, 40, 30, and 10 μm cell strainers) and centrifuged at 300 × g for 10 min at 4°C. Cells were resuspended in 10 mL medium (DMEM/F-12 + 10% FBS + P/S) supplemented with biotinylated antibodies, rat anti-mouse CD16/32 (0.65 μg/million cells), rat anti-mouse CD45 (1.5 μg/million cells), and rat anti-mouse Ter119 (5 μg) and incubated for 30 min at 37°C on an incubator shaker at 60 rpm. Resuspended cells were then incubated with pre-washed streptavidin-coated magnetic particles for 30 min at room temperature to remove undesired cells. Selected cells were then resuspended in 10 mL medium (DMEM/F-12 + 10% FBS + P/S) and incubated for 30 min at 37°C in a sterile Petri dish to allow residual fibroblasts to adhere to the bottom of the dish. Suspended cells were transferred in plates pre-coated with mouse IgG for 2 h in a 5% CO_2_ incubator to remove macrophages. Unattached cells were collected and centrifuged at 300 × g for 10 min at 4°C. Cell pellets were then resuspended in a complete mouse medium (CMM: DMEM/F-12 supplemented with 2 mM L-glutamine, 0.25% bovine serum albumin, 10 mM HEPES, 0.1 mM non-essential amino acids, 0.05% ITS, 100 μg/mL primocin, and 10% newborn calf serum). The viability of harvested AT2 cells was assessed by the trypan blue exclusion assay followed by cell counting for the yield.

The purity of isolated AT2 cells was confirmed by the Papanicolaou stain, immunofluorescent staining with anti-proSP-C antibody, and fluorescence-activated cell sorting (FACS) with anti-EpCAM antibody. 1) **PAP stain**: freshly isolated AT2 cells suspended in DPBS + 10% FBS (2 × 10^5^ cells/mL) were centrifuged at 600 rpm for 4 min by a Shandon CytoCentrifuge. Slides were air-dried overnight and stained using a modified PAP stain method^2^. Briefly, slides were stained with hematoxylin for 3.5 min. After rinsing in dH_2_O, slides were incubated in lithium carbonate for 2 min and rinsed with dH_2_O again. Slides were dehydrated in serial ethanol dilutions, then in xylene: ethanol (1:1) for 30 s, and finally in 100% xylene for 60 s. Slides were mounted with the Permount mounting medium. Randomly selected images were captured from 6 independent experiments with an Olympus BX41 microscope (40 ×). AT2 cells with large nuclei and blue colored granules spread in the cytoplasm were counted and calculated for purity (%). 2) **Immunofluorescent stain**: cytospanned freshly isolated AT2 cells and cells cultured on coverslips for 48 h were incubated with rabbit anti-proSP-C (1:500) overnight at 4°C. Secondary antibody, either Alexa Flour 488 or 568-conjugated anti-rabbit IgG, was added to recognize proSP-C antibody. 3) **FACS**: Briefly, cells were resuspended in staining buffer containing 0.1 mM EDTA. Cells were incubated with Alexa Fluor 488 conjugated anti-mouse CD326 for 30 min on ice in the dark. Cells were washed and resuspended in staining buffer and analyzed for antibody expression on BD FACSCalibur^™^. Data were analyzed with FlowJo 10.1 software. The purity of the final cell suspensions was about 93% (**Fig. S1A-C**). Cell viability was approximately 94%. Further, FACS showed that approximately 95% of cells were EpCAM positive (**Fig. S1D-E**).

#### Human AT2 isolation

Human epithelial AT2 cells were isolated from six human lungs not used for transplantation by the Northern California Transplant Donor Network as previously described^3,4^. Three of the lungs were normal and three of the lungs met ARDS criteria before they were harvested. After cold preservation at 4°C, the right middle lobe was selected for cell isolation if no apparent signs of consolidation or hemorrhage by gross inspection were seen. The pulmonary vasculature was first flushed antegrade and retrograde with PBS to remove the remaining blood from the microcirculation. The distal airspaces were then lavaged 10 times with Ca^2+^, Mg^2+^-free PBS solution (37°C) containing 0.5 mM EGTA and 0.5 mM EDTA. Elastase, 13 U/mL in Ca^2+^, Mg^2+^-free Hanks’ balanced salt solution, was instilled into the distal airspaces by segmental bronchial intubation. The lobe was then digested at 37°C for 45 – 60 min. After digestion, the lobe was further minced in the presence of FBS and DNase I (500 μg/mL). The cell-rich fraction was filtered sequentially through multiple layers of sterile gauze and 100- and 20- μm nylon meshes (Spectra/Mesh). The filtrate was further layered onto a discontinuous Percoll (Sigma) density gradient of 1.04 and 1.09 g/mL solution and centrifuged at 400 × *g* for 20 min. The interface containing both AT2 cells and alveolar macrophages was collected and further centrifuged at 800 rpm for 10 min, and the cell pellet was washed and resuspended in Ca^2+^, Mg^2+^-free PBS containing 0.5% FBS. The cells were incubated with magnetic beads coated with anti-CD 14 antibodies (Dynabeads^®^ CD14, Invitrogen) at 4 C for 40 min, and the majority of the macrophages were then selectively depleted with a Dynal magnet (Dynal Biotech, Oslo, Norway). The cell suspension was further incubated on Petri dishes coated with human IgG antibodies (Sigma) overnight at 37°C to remove the remaining macrophages and fibroblasts. Cell viability was assessed by the trypan blue exclusion method. AT2 cell purity was evaluated by Papanicolaou (Pap) staining.

#### CD44^+^ AT2 cells sorting and analysis by FACS

Freshly isolated cells were seeded on either collagen IV (for mouse AT2, 10 μg/cm^2^) or collagen I (for human AT2, 10 μg/cm^2^) coated plates for 24 – 36 h to revive CD44 expression diminished by digestive enzymes. Both unattached and attached (trypsinized) cells were collected and blocked with 1% BSA, 4% normal goat serum in PBS. Cells were stained with AF488-EpCAM (BioLegend), APC anti-human CD44 (BioLegend), and their respective isotypes. Cells were sorted using a Beckman Coulter MoFlo high-speed cell sorter. Unstained, isotype and single-color controls were performed. The gates for CD44 and EpCAM were set based on the results of isotype, and single-color controls were run in parallel. The results were analyzed using FlowJo 10.1 software.

#### 3D polarized AT2 monolayers and geometrical measures

Transwell inserts (Costar 3470: 0.4 μm pore size, 0.33 cm^2^ area; Corning Costar, USA) were pre-coated with mouse laminin 1 at 10 μg/cm^2^ (for mouse AT2 cells; Trevigen, USA) for 4 – 6 h at 37°C or with rat tail collagen I at 10 μg/cm^2^ (for human AT2 cells; Trevigen, USA) for 1 h at 37°C. Freshly isolated AT2 cells were seeded at 10^6^ cells/cm^2^. The CMM medium (600 μL) was added to the basolateral side of each transwell. The culture medium on the basolateral side was replaced with a serum-free medium in 48 h post seeding. Transepithelial resistance (R_T_, Ω) and potential difference (V_T_, mV) were measured using an epithelial voltohmmeter (EVOM: World Precision Instrument, USA) in 72 h. The culture medium was then replaced with serum-free media and maintained with the air-liquid interface until day 5. Monolayers were maintained in a humidified 5% CO_2_ air incubator at 37°C. Stacked image files (avi format) were segmented with 3D Objects Counter. Surface files were used for both 3D geometrical and shape measures. The 3D surface plot was used to generate surface graphs. This 3D model has the following advantages: 1) Stromal cells are not used (feeder-free). 2) Tight junction proteins are generated to develop an epithelial barrier in an *in vivo* pattern. 3) Cells are polarized to separate apical and basolateral membranes. 4) Cytosolic and cell surface signal proteins are expressed and distributed in a polarized way. 5) It is suitable for testing the permeability of the epithelial barrier. 6) It complimentarily overcomes the limitations of 3D organoids for studying barrier function and polarization of signaling proteins.

#### Measurements of bioelectric properties in AT2 monolayers

Transepithelial short-circuit current (I_SC_, μA/cm^2^) in AT2 monolayers was measured with an 8-channel voltage-clamp amplifier (Physiological instruments, USA) as previously described^5^. Briefly, AT2 monolayers were mounted in the vertical Ussing Chambers bathed with solutions containing (in mM): 120 NaCl, 25 NaHCO_3_, 3.3 KH_2_PO_4_, 0.83 K_2_HPO_4_, 1.2 CaCl_2_, 1.2 MgCl_2_, 10 HEPES, 10 mannitol (apical compartment), or 10 D-glucose (basolateral compartment). Each solution was iso-osmotic. The transwell cultures were bubbled continuously with a gas mixture of 95% O_2_-5% CO_2_. The transmonolayer potential was short-circuited to 0 mV, and a 10-mV pulse of 1-s duration was imposed every 10 s to monitor transepithelial resistance. Data were collected with the Acquire and Analyze program (version 2.3; Physiologic Instruments). When the Isc level reached a plateau, compounds were pipetted to the apical compartment.

#### 3D organotypic cultures of AT2 cells

AT2 cells were cultured as organoids as previously described^6^. Briefly, Mlg-2908 cells (2 × 10^5^ cells/mL) were mixed with 6,000 primary AT2 cells and pelleted down for each transwell. Cells were then resuspended into a 100 μL mixture (1:1) of growth factor reduced matrigel (Corning, USA) and organoid medium (DMEM/F12 supplemented with 2 mM L-glutamine, 10 % active FBS, 1% ITS, and 10 μM ALK inhibitor). On each 0.33 cm^2^ insert (Corning Costar, USA) 50 uL mixed cells were seeded and incubated at 37°C for 30 min to allow the matrix to solidify. Then 410 μL culture medium was added to the bottom well and changed half (200 μL) of the medium every other day. The DIC images of organoids (diameter ≥ 50 μm) were visualized with an Olympus IX 73 microscope (4× objective, Olympus, Japan) with an Hamamatsu photonics CMOS camera (Orca Flash 4.0; 2,048 × 2,048 pixels) on a designed day post seeding. The surface area of individual organoid was measured with Image J. The sum of all organoids on each transwell insert was the total surface area. Histologic images of organoids were captured from sliced matrigel. Matrigel containing organoids was fixed with 4% paraformaldehyde in PBS for 1 h, dehydrated with increasing ethanol grades, and embedded in paraffin blocks. Sections at the thickness of 7 μm were fixed for an additional 5 min in 4% paraformaldehyde, then H & E stain was performed as described previously^7^.

#### Quantification of AT1 and AT2 cells in organoids and monolayers

Anti-pdpn and anti-sftpc antibodies were used to detect AT1 and AT2 cells, respectively. Fluorescence conjugated secondary antibodies, goat anti-hamster AF568 and goat anti-rabbit IgG AF488 were used. Fluorescent images were projected with a Zeiss LSM 510 confocal microscope and stacked with a Fiji plug-in for ImageJ. Monolayers and organoids were scanned for Z sections with optimal depth from top to bottom. Images were stacked for pdpn and sftpc signal separately to count the number of positive cells precisely with a cell counter plug-in of ImageJ. Alternatively, cells were collected from organoids and monolayers and sorted by FACS. Each slide was scanned for at least 6 different fields (n = 3 animals/experiments). For 3D organoids, all Z sections were stacked from top to bottom and saved as .avi files.

#### EdU assay for DNA synthesis

AT2 cells with active DNA synthesis in organoids and monolayers were detected with a Click-iT^™^ EdU assay kit. Organoids and monolayers from both wt and *Plau^−/−^* groups were stained with the Click-iT^™^ EdU Alexa flour 488 following the manufacturer’s instructions. Images were captured and analyzed for the percentage of EdU^+^ cells in different experimental groups. Ten randomly selected images across the monolayer from 3 independent experiments were captured and counted for total cells and EdU^+^ portion. For organotypic cultures, all 3 organoid types were scanned (n = 5 colonies for each type). The z sections were stacked separately for DAPI (blue) and EdU (green) to produce a 3D structure of organoids for counting total and EdU^+^ cells, respectively, using a cell counter plug-in for the ImageJ. The percentage of EdU^+^ cells was calculated for each group, and the difference among groups was compared statistically.

#### Application of A6 peptide and CD44-blocking antibody to organoids

A6 and scrambled A6 (sA6) peptides were synthesized by Genscript. Stock solutions (2 mM) were prepared by dissolving peptides in water according to the manufacturer’s instruction. For 3D matrigel cultures, sorted *Plau^−/−^* AT2 cells were preincubated with either A6 or sA6 peptide (1 μM) for 30 min at room temperature. Wt AT2 cells were treated with 30 ng/mL CD44-blocking antibody for 30 min at room temperature. The same concentrations of A6, sA6 peptides, and CD44-blocking antibody were added to both the matrigel/medium mix and the culture medium placed under the transwell inserts. The culture medium was changed every 48 h.

#### Organoid dissociation and FACS

For analysis of AT2 cell proliferation and differentiation, AT2 organoids from different experimental groups were isolated from matrigel with dispase (10 U/mL) and dissociated in 0.25% trypsin-EDTA to get a single-cell suspension. Cells were then stained with antibodies AF488 conjugated EpCAM, APC conjugated ICAM, and APC conjugated PDPN. Gates for both colors were set by unstained cells and isotype controls for each antibody. We have used a strategy to use double color staining to enhance the separation of AT1 and AT2 cells in the cell suspension of organoids. Cells were analyzed by FACSCaliber^™^ (BD, USA), and the results were analyzed using FlowJow 10.1 software.

#### Statistical analysis

Data were presented as mean ± s.e.m. No animals were excluded. Normality tests were performed to determine whether the data were parametric or not. If the data were normally distributed and the variance between groups was not significantly different, mean differences in measured variables between the experimental and control group were assessed with the Student’s two-tailed t-tests or one-way ANOVA followed by the Tukey’s or Bonferroni’s post hoc test. Otherwise, the Mann-Whitney U test was applied for analyzing non-parametric results. Two-way ANOVA followed by Sidak’s multiple-comparison test was used for multiple comparisons. Meanwhile, the actual power of the sample size was analyzed. Mean differences were considered statistically significant at the levels of P < 0.05, P < 0.01 and P < 0.001. Origin Pro 2018 was used for statistical analysis and plotting.

**Figure S1.**
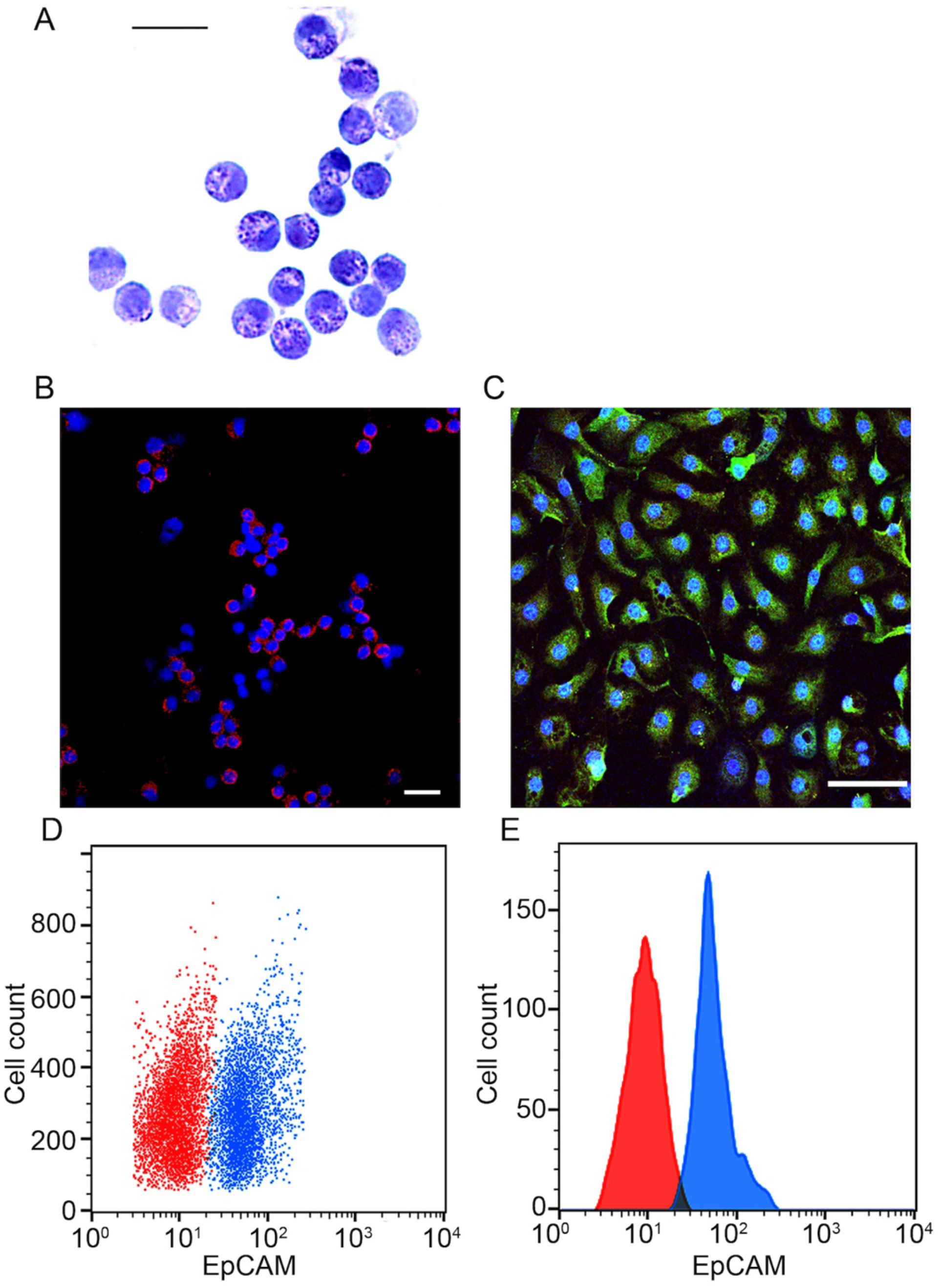
Cytological characterization of mouse alveolar type 2 (AT2) epithelial cells. The viability of isolated AT2 cell preparations was examined with trypan blue assay, then for its purity with pap staining, immunostaining with anti-sftpc antibody, and FACS analysis. A. Representative AT2 cell image of pap stain showing featured cytoplasmic granules. A total of 1,000 cells counted to calculate the purity. B-C. Representative immunostaining images of cytospanned (B) and cultured AT2 cells post 72 h (C), respectively, probed with an anti-pro-surfactant protein C (pro-SPC) antibody. Nuclei were counterstained with Hoechst dye (blue). D-E. The purity of freshly harvested mouse AT2 cells was analyzed with FACS. Cells were incubated with an anti-EpCAM antibody. Scale bar in B, C, and D is 100 μm. Data in a is mean ± sem and analyzed with the Student t-test. ***indicates P < 0.001 vs wild type (wt) controls. (Files: 08212017 D12 Phase Contrast 4X. uPA vs WT organoids for CFE Day8 and Day12, origin pro file).

**Figure S2.**
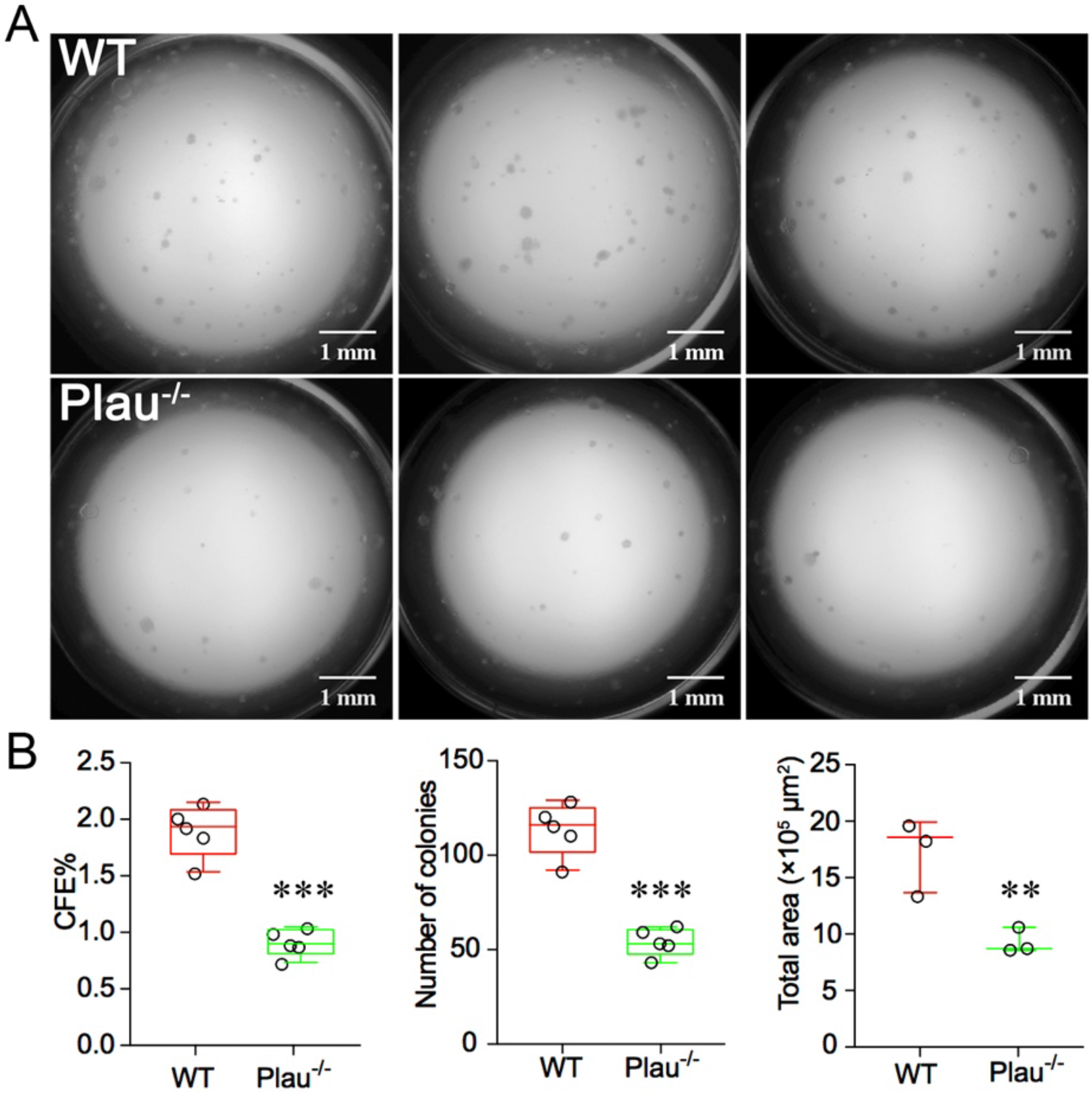
Comparison of AT2 organoids between wt and *Plau^−/−^* groups 12 days post planting in 3D matrigel. A. DIC images. Phase-contrast images were captured (4 ×). Scale bar, 1 mm. B. Colony-forming efficiency (CFE, left), counting (middle), and surface area (right). Colonies with a diameter of < 50 μm were not counted. *** P < 0.001 and **P < 0.01 vs controls. n = 7. (File: 08212017 D12 Phase Contrast 4 ×; project interim summary of Nov 2017.key. uPA vs WT organoids for CFE Day8 and Day12.opj.)

**Figure S3.**
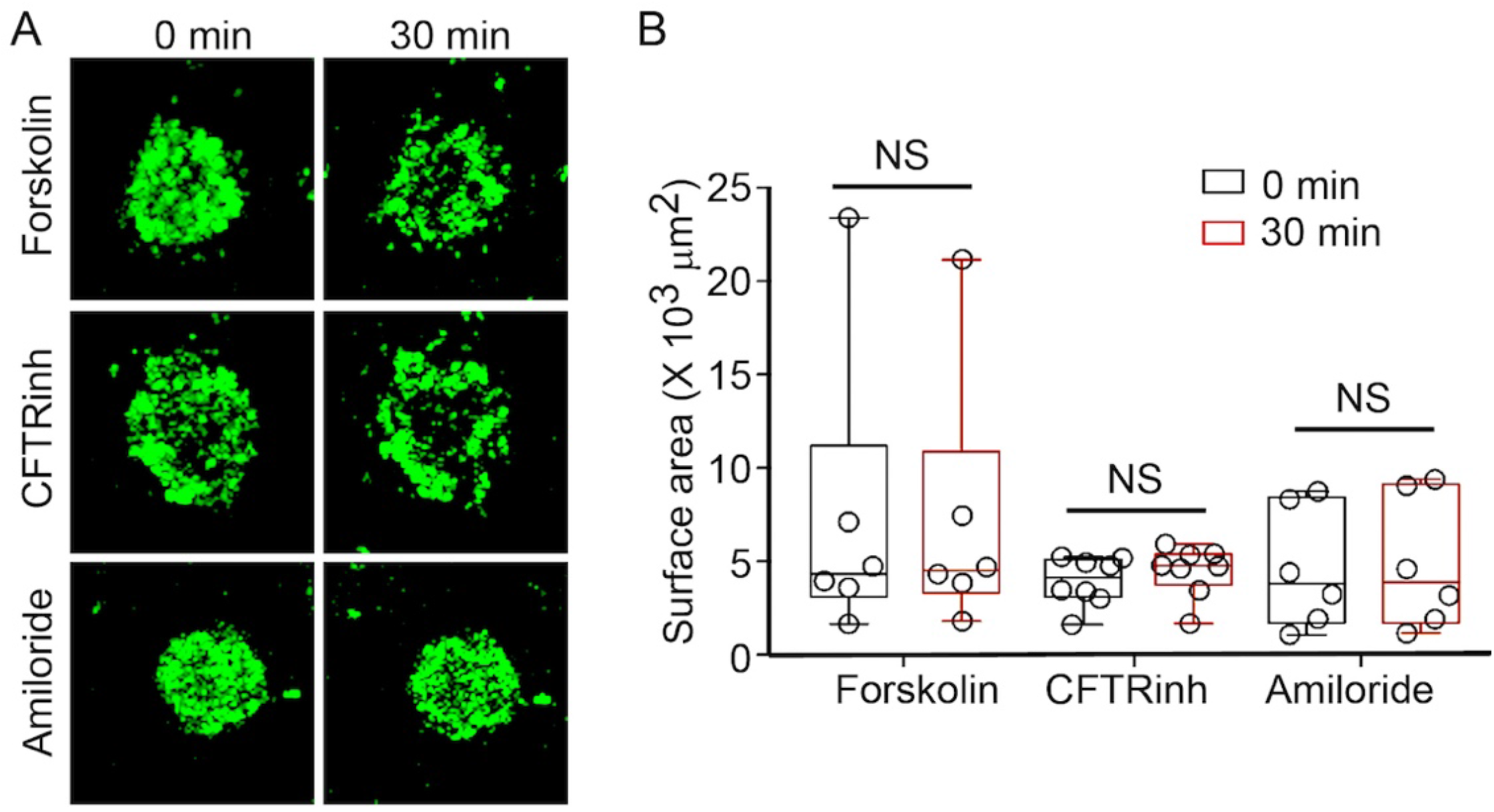
Measurement of organoid permeability. A. Organoid images with live-cell cytotracker calcien green at 0 (left) and 30 min (right) after addition of forskolin (10 μM, CFTRinh 20 μM, and amiloride 100 μM). Full length, 200 μm. B. Surface area of organoids. n = 8. NS, not significant. (File: AT2 D7 organoid 100um forskolin calcien green. Normalized area of AT2 organoids treated with forskolin amiloride and CFTR.ppt.)

**Figure S4.**
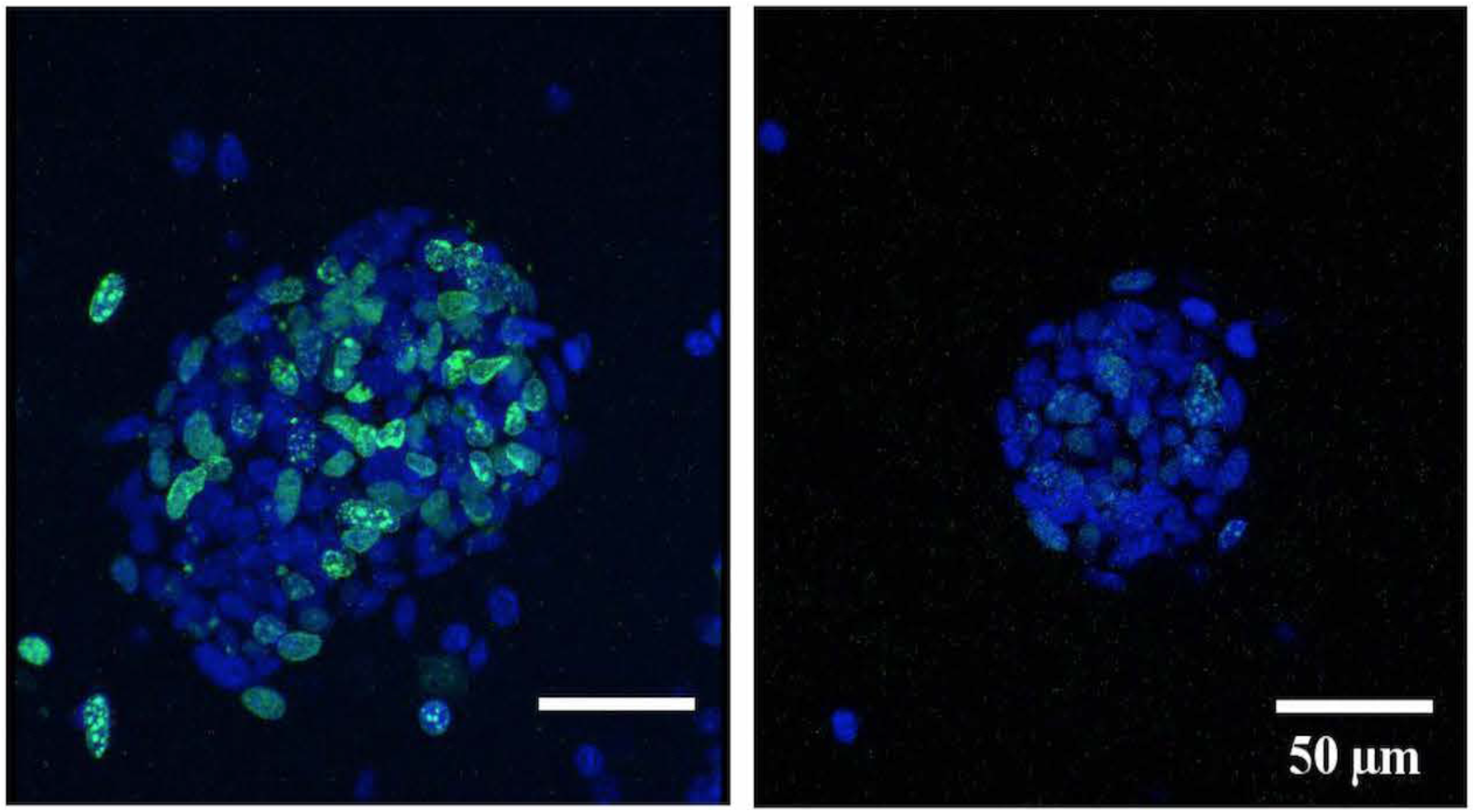
3D video clips of organoids stained with EdU. Representative organoids of wt (left) and *Plau^−/−^* (right) AT2 cells were imaged (x-y), stacked of z sections, and visualized by the ImageJ. EdU^+^ cells were stained as green nuclei. The volume was 9,111 and 2,806 voxels for wt and *Plau^−/−^* organoids, respectively. The geometrical surface area was 41,380 (wt) and 12,828 pixels (*Plau^−/−^*). n = 6. (File name: 12222017 EdU Staining D6 organoid wt vs plau ko. Avi or tif files.)

**Figure S5.**
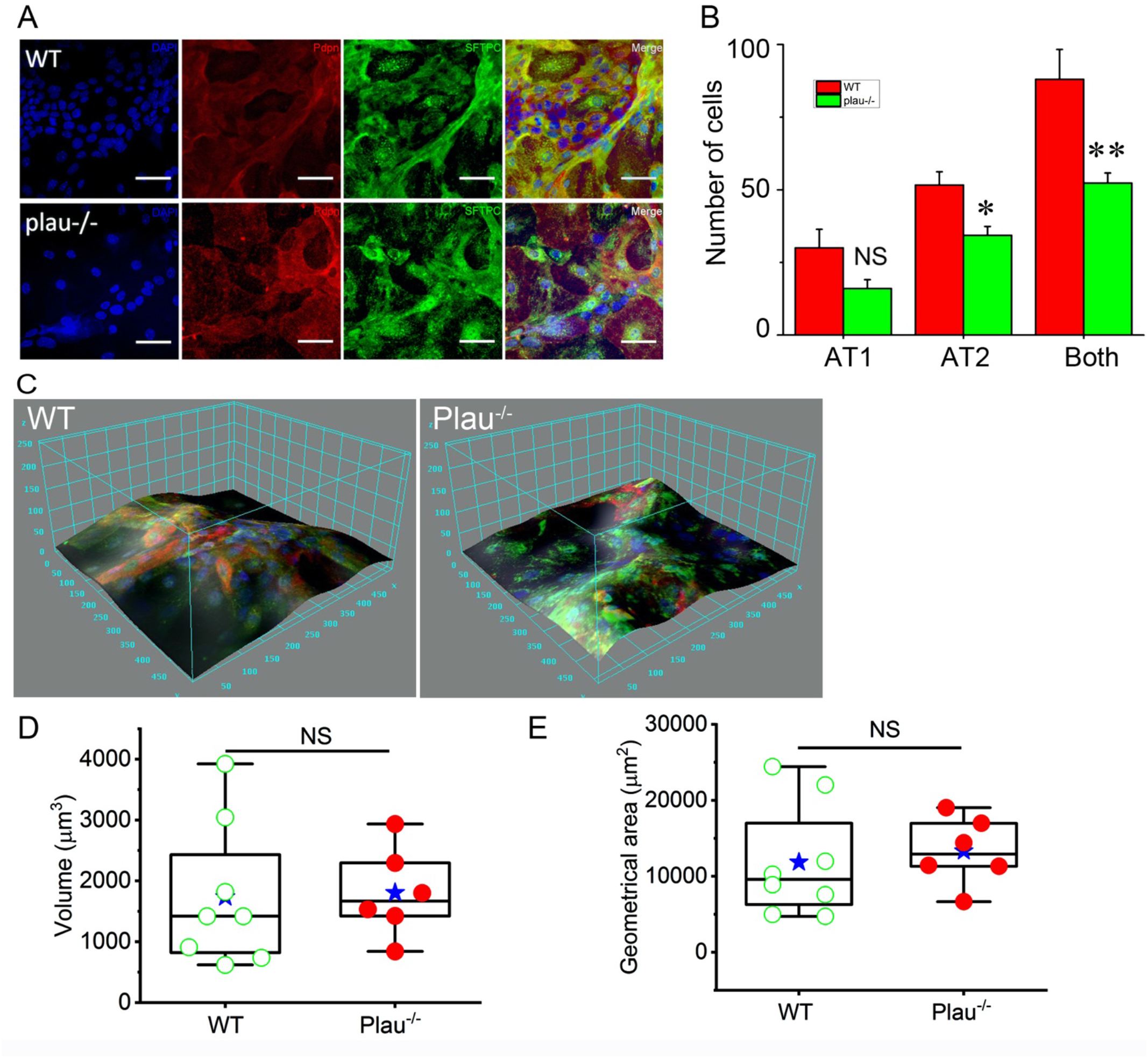
Effects of A6 peptide and CD44 blocking antibody on the formation of AT2 organoids. A. Representative DIC images of AT2 organoids. From left to right, wt control organoids (WT), wt organoids treated with CD44 blocking antibody (CD44 Ab), *Plau^−/−^* AT2 organoids (*Plau^−/−^*), and *Plau^−/−^* organoids treated with A6 peptide (A6). 4 ×. B. Organoid number. n = 6 replicates per experiment, n = 6 mice/genotype. * P < 0.05 and **P < 0.01 vs controls. C. Surface area of organoids. * P < 0.05 and **P < 0.01 vs controls. n=6. Data in B & C were mean ±sem and analyzed by Student’s t-test. (File names: Supp fig for fig 7.jpg.)

**Figure S6.**
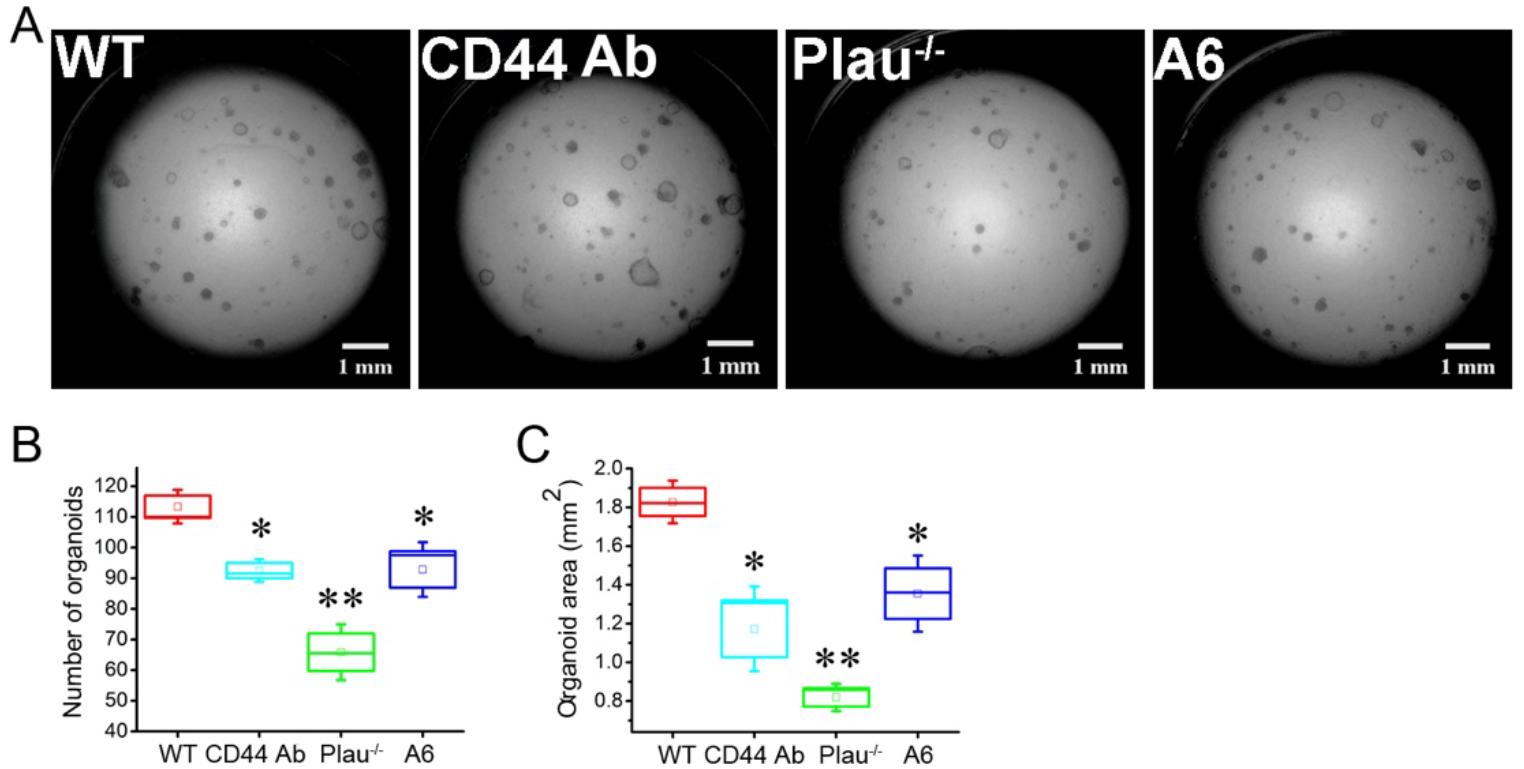
Analysis of polarized AT2 monolayers. A. Visualization of polarized tight mouse AT2 monolayers for wt (top) and *Plau^−/−^* (bottom) mice. From left to right were images of nuclei (DAPI), AT1 cells (Pdpn), AT2 cells (Sftpc), and merged images. Scale bar, 50 μm. B. Comparison of AT1 and AT2 cells between wt and *Plau^−/−^* monolayers. n = 6. * P < 0.05 and ** P < 0.01 vs wt controls. C. 3D surface plots. D-E. 3D geometrical measures for volume (D) and surface area (E). n = 5 for wt and n = 8 for *Plau^−/−^* group. NS, no significant. (File name: Figure S7 folder).

